# Tree diversity increases carbon stocks and fluxes above- but not belowground in a tropical forest experiment

**DOI:** 10.1101/2024.06.20.599915

**Authors:** Florian Schnabel, Joannès Guillemot, Kathryn E. Barry, Melanie Brunn, Simone Cesarz, Nico Eisenhauer, Tobias Gebauer, Nathaly R. Guerrero-Ramirez, I. Tanya Handa, Chris Madsen, Lady Mancilla, Jose Monteza, Tim Moore, Yvonne Oelmann, Michael Scherer-Lorenzen, Luitgard Schwendenmann, Audrey Wagner, Christian Wirth, Catherine Potvin

## Abstract

International commitments advocate large-scale forest restoration as a nature-based solution to climate change mitigation through carbon (C) sequestration. Mounting evidence suggests that mixed compared to monospecific planted forests may sequester more C, exhibit lower susceptibility to climate extremes and offer a broader range of ecosystem services. However, experimental studies comprehensively examining the control of tree diversity on multiple C stocks and fluxes above- and belowground are lacking. To address this gap, we leverage data from the Sardinilla experiment in Panama, the oldest tropical tree diversity experiment which features a gradient of one-, two-, three- and five-species mixtures of native tree species. Over 16 years, we measured multiple above- and belowground C stocks and fluxes, ranging from tree aboveground C, over leaf litter C production, to soil organic carbon (SOC). We show that tree diversity significantly increased aboveground C stocks and fluxes, with a 57% higher gain in aboveground tree C in five-species mixtures compared to monocultures (35.7±1.8 *vs* 22.8±3.4 Mg C ha^-1^) 16 years after planting. In contrast, we observed a net reduction in SOC (on average -11.2±1.1 Mg C ha^-1^) and no significant difference in SOC_3_ stocks (the predominantly tree-derived, i.e., C_3_ plant-derived SOC fraction) between five-species mixtures and monocultures (13.0±0.9 *vs* 15.1±1.3 Mg C ha^-1^). Positive tree diversity effects persisted despite repeated climate extremes and strengthened over time for aboveground tree growth. Structural equation models showed that higher tree growth in mixtures enhanced leaf litter and coarse woody debris C fluxes to the soil, resulting in a tightly linked C cycle aboveground. However, we did not observe significant links between above- and belowground C stocks and fluxes. Our study elucidates the mechanisms through which higher tree diversity bolsters the climate mitigation potential of tropical forest restoration. Restoration schemes should prioritize mixed over monospecific planted forests.

## Introduction

Forest restoration is promoted as a key strategy for mitigating climate change through carbon (C) sequestration. Several global initiatives, such as the Bonn Challenge and the New York Declaration on forests, aim to restore 350 Mha of forests by 2030. Forest restoration, in particular in the tropics, has the most significant climate mitigation potential of 20 proposed nature-based solutions, potentially sequestering up to 10.1 PgCO_2_ equivalents per year (Griscom et al., 2017). However, massive reforestation efforts should ensure the protection of land for agriculture (Dooley et al., 2022) and avoid the replacement of other ecosystems, such as natural grasslands (Parr et al., 2024; Seddon, 2022). A solution is to target the vast areas of degraded land suited for forest growth (Bauhus et al., 2010). Current reforestation pledges largely focus on monospecific planted forests often with non-native tree species (Lewis et al., 2019) despite the mounting evidence that tree species-diverse planted forests (hereafter mixed planted forests or mixtures) can exhibit lower susceptibility to stress and disturbances such as droughts and storms while simultaneously providing a broader range of ecosystem services such as C sequestration and storage, biodiversity conservation and cultural services at higher levels than monospecific plantations (Messier et al., 2021). Consequently, mixed planted forests, particularly if established with native tree species, better fulfil current international targets such as the Kunming-Montréal Global Biodiversity Framework (CBD, 2022).

Biodiversity-ecosystem function theory suggests that mixed planted forests may outperform monocultures in terms of productivity, through complementary resource partitioning across species, abiotic facilitation or biotic feedbacks (Barry et al., 2019). Thus, mixed planted forests may also outperform monocultures with respect to their role in climate regulation. Indeed, there is accumulating evidence that mixed planted forests can sequester more C above- and belowground than their monoculture counterparts (Chen et al., 2023; Lecina-Diaz et al., 2018; Messier et al., 2021; van der Sande et al., 2017; Warner et al., 2023; Xu et al., 2020). More importantly, mixtures may also be more stable than monocultures in the face of climate extremes or climate variability in general (Isbell et al., 2018), as some species may ‘insure’ the community against the reduced functioning of other species (Yachi and Loreau, 1999). Mixed planted forests indeed feature higher temporal stability of biomass production than monocultures during periods with variable climatic conditions, including particularly wet and dry years (Jucker et al., 2014; Schnabel et al., 2021). However, most existing studies assessed (or indirectly inferred) tree diversity effects on C stocks, fluxes, and stability in terms of aboveground tree C (AGC), with fewer studies examining root C or soil organic C (SOC) (e.g. Xu et al., 2020) and even fewer ones the C fluxes above- and belowground, connecting these C pools.

We posit that tree species richness (hereafter tree diversity) may affect C stocks and fluxes both above- and belowground. C stocks refer to the C stored in reservoirs such as AGC or SOC, whereas C fluxes are the flow of C between these reservoirs over time. We anticipate comparable diversity effects above- and belowground as has been shown in grassland experiments (Ravenek et al., 2014; Weisser et al., 2017). Indeed, there is evidence for significant tree diversity effects on various C fluxes, ranging from enhanced leaf litter (Huang et al., 2017) and coarse woody debris (CWD) production (Liu et al., 2018) to enhanced microbial respiration and thus decomposition (Chen et al., 2020). Ultimately, the balance of these different C fluxes determines net tree diversity effects on C sequestration in forests (Liu et al., 2018). For example, tree diversity is often reported to increase tree biomass production (hence C gain, Potvin et al., 2008) but may, in some cases, also increase tree mortality (hence C loss, Searle et al., 2022). Similarly, tree diversity may increase SOC through diversity-induced enhancements of C inputs into the soil via plant litter or root exudate production, but changes in soil community and functioning may also enhance C losses due to decomposition (Chen et al., 2020; Handa et al., 2014; Lange et al., 2015). Due to these complex interactions, no net effect of tree diversity on SOC was reported in some studies (e.g. Martin-Guay et al., 2022).

Comprehensive assessments of the multiple C stocks and fluxes in forests and their intricate relationships are scarce (Xu et al., 2020). A notable exception is Liu et al. (2018), who studied naturally established subtropical forests in China, revealing significant positive effects of tree diversity on AGC, root C, CWD, and SOC, and significant correlations between tree diversity and AGC, CWD, and leaf litter production. However, in complex natural environments like forests, environmental variation and tree diversity interactively influence carbon stocks and fluxes (van der Sande et al., 2017). Despite attempts using structural equation models (SEMs) to identify direct and indirect relationships (Chen et al., 2018; Chen et al., 2023; Li et al., 2020), mechanistically disentangling these drivers remains challenging in observational studies. Planted tree diversity experiments, which were specifically designed to compare monocultures and mixtures of increasing diversity while controlling for environmental variation and holding tree density constant (Depauw et al., 2024; Scherer-Lorenzen et al., 2005), offer an ideal setting for elucidating linkages among C stocks and fluxes. Until recently, the young age of most tree diversity experiments, the slow development of trees in boreal and temperate experiments and the different response times and dynamics of C compartments over the course of stand development (e.g. faster responses of aboveground compared to belowground C; Ravenek et al., 2014) prevented analysing the temporal dynamics of tree diversity effects on C stocks and fluxes. Moreover, a temporal perspective on C residence time, i.e. the time C is stored within a reservoir, is a prerequisite for assessing the stability of C storage under climate variability and investigating whether tree diversity’s control on C stocks and fluxes increases as forest stands develop. Increases in ecosystem functioning over time in more diverse tree communities have been demonstrated for aboveground tree productivity (Guerrero-Ramírez et al., 2017; Jucker et al., 2020) but not for multiple C stocks and fluxes and their relationships.

Here, we use data on temporal changes in ten C-related stocks and fluxes measured in the oldest tropical tree diversity experiment, the Sardinilla experiment established in 2001 in Panama, which is part of the global network of tree diversity experiments (TreeDivNet). After two decades of C-related research (Cesarz et al., 2022; Coll et al., 2008; Guerrero-Ramírez et al., 2016; Guillemot et al., 2020; Hutchison et al., 2018; Kunert et al., 2019; Kunert et al., 2022; Madsen et al., 2020; Moore et al., 2018; Murphy et al., 2008; Potvin et al., 2011; Ruiz-Jaen and Potvin, 2011; Sapijanskas et al., 2013; Sapijanskas et al., 2014; Scherer-Lorenzen et al., 2007; Schnabel et al., 2019; Wolf et al., 2011) and due to the comparably fast tree growth in the tropics, the Sardinilla experiment features a wealth of C-related variables above- and belowground for planted forests with 1–5 tree species and considerably large-sized trees (with the tallest trees over 25 m), which we leverage here to explore tree diversity effects on C stocks and fluxes across 16 years (2001– 2017). Since planting, the experiment experienced repeated climate extremes including a severe El Niño-driven drought and a Hurricane (see methods). Although it is not possible to disentangle the intertwined impacts of stand development and these climate extremes, their occurrence has provided us with the unique opportunity to evaluate the role of tree diversity for C stocks and fluxes in the face of severe climate events. We anticipate a stronger positive tree diversity effect at later stages of stand development, due to enhanced ecosystem functioning in more diverse tree communities over time (Guerrero-Ramírez et al., 2017) and a higher stability of diverse communities to climatic extremes (Schnabel et al., 2021). Specifically, we tested the following hypotheses: (H1) C stocks and fluxes increase with increasing tree diversity. (H2) Positive tree diversity effects on C stocks and fluxes increase with stand development despite repeated climate extremes. Finally, we use SEMs to test how C stocks, fluxes, and their control through tree diversity are connected through direct and indirect relationships above- and belowground using 12 explicit hypotheses (Table S1).

## Methods

### Description of the study site

This study is based on data collected over 16 years in the Sardinilla planted forest. Established in 2001 (Scherer-Lorenzen et al., 2005) Sardinilla is the oldest tropical experiment of the International Network of Tree Diversity Experiments (TreeDivNet; https://treedivnet.ugent.be/, Verheyen et al. (2016)). The site was planted with six native tree species on a former pasture dominated by C_4_ grasses without trees, namely *Luehea seemannii* Triana & Planch (Ls), *Cordia alliodora* (Ruiz & Pavon) Oken (Ca), *Anacardium excelsum* (Bert. & Balb. Ex Kunth) Skeels (Ae), *Hura crepitans* L. (Hc), *Tabebuia rosea* (Bertol.) DC. (Tr) and *Cedrela odorata* L. (Co). Species were chosen based on their relative growth rates in natural forests of the region, always combining fast (Ls, Ca), intermediate (Ae, Hc), and slow (Tr, Co) growing species in mixtures to promote divergence in traits and shade tolerances (Scherer-Lorenzen et al., 2005). A total of 24 plots ranging from 0.2025 to 0.2304 ha (approximately 45 × 45 m) were established featuring 12 monocultures (2 plots per species), six three-species mixtures with each species present in two plots, and six plots with all tree species. Diversity treatments were randomly allocated to plots. Trees were planted at a constant density of 3 × 3 m following standard reforestation practices in the region. Due to high mortality experienced by Ca in the two years after planting, only 22 plots were maintained over the 16 years of the experiment. This paper thus considers the effect of three diversity levels, grouped as 1, 2, 3, and 5 species. Elevation across the site ranges from a ridge at 79 m ASL to low areas at 67 m ASL (Healy et al., 2008) resulting in a gradient of soil types ranging from Vertic Luvisol on the ridge to Gleyic Luvisol in the low part of the plantation (Oelmann et al., 2010). The average pH of the top 10 cm of the soils was 4.8 in both 2001 and 2011 (Moore et al., 2018). An average clay content of 65%, a high cation exchange capacity and base saturation and the underlying carbonate-rich parent material contribute to a high nutrient availability (Oelmann et al., 2010). Further details on the Sardinilla tree diversity experiment can be found in Scherer-Lorenzen et al. (2005) and Potvin and Dutilleul (2009).

We examined three periods: an early (p1), a mid (p2), and a late period (p3) of plantation development (Fig. S1). These periods were characterized by repeated climate extremes, with the mid-period featuring an extremely wet year (2010) and the late period a severe El Niño-driven hotter drought (2015) triggering growth reductions and elevated tree mortality (Browne et al., 2021; Detto et al., 2018; Hutchison et al., 2018; Schnabel et al., 2019). Subsequently, in November 2016, the experiment was hit by Hurricane Otto, a tropical storm that formed off the coast of Panama in the Caribbean Sea inducing stem breakages in the experiment. Climatic conditions at the Sardinilla experiment were characterized in terms of annual mean temperature, precipitation sum, and drought index (Standardized Precipitation Evapotranspiration Index (SPEI; Vicente-Serrano et al., 2010)), with all climate variables illustrating the climate extremes described above in terms of temperature and precipitation extremes and drought conditions (Fig. S2).

### Data collection

We measured ten compartments of the forest C cycle, namely aboveground tree C (AGC), tree coarse root C (CRC), coarse woody debris C (CWDC), C in herbaceous biomass (herbaceousC), leaf litter C production (litterC), soil organic C (SOC), leaf litter decomposition, root decomposition, soil microbial biomass C (C_mic_), and soil respiration. In addition, we included canopy opening as a co-variable, with potentially important influences on C stocks and fluxes. Each variable was measured during three periods of plantation development: an early (2001 for SOC and 2005, 2006 & 2007 for the other variables), a mid (2011, 2012 & 2013), and a late period (2016 & 2017) (see Fig. S1 for a timeline). Three variables were only measured in some periods: C_mic_ in the mid (2013) and late (2017), root decomposition in the mid (2012), and soil respiration in the late (2017) period. If not stated otherwise, the sample size for all variables was n = 22 plots. We aggregated variables to periods in our analysis as not all variables were measured in all years. To scale individual tree measurements up to community measures, we used diameter and height inventories of all trees in the plantation conducted at the end of each growing season (December-January) in 2005, 2012, and 2016. The measurement of individual variables is described briefly below, with details provided in the Supplementary Methods.

#### Aboveground tree carbon

Aboveground tree biomass (AGB) estimates were based on species- and diversity-specific allometric equations developed after harvesting and measuring 150 and 167 trees in the experiment in 2005 and 2017. AGB was calculated as the sum of trunk and branch biomass (excluding leaves to focus on the more permanent C-components of the trees). Allometric models are provided in Supplementary Methods. The best-fitting models were then combined with annual diameter and height inventories of all trees in the experiment conducted at the end of each growing season (December-January) to estimate the AGB of each tree in each period. Allometric models calibrated in 2017 were used for the mid and late period. AGB was converted to aboveground tree C (AGC) using species-specific trunk C concentrations (Elias and Potvin, 2003).

#### Coarse root carbon

To estimate coarse root C (CRC) we relied on root:shoot ratios based on two different root excavation campaigns in the experiment. For CRC in the early period, we relied on root:shoot ratios developed in 2004 from excavating of three-year-old trees, where ratios were obtained for Ls, Co, and Hc, and mean values were used for Ae and Tr (Coll et al., 2008). For CRC values in the mid and late period, we used species-specific root:shoot ratios developed in 2017 (Guillemot et al., 2020). The species-specific root:shoot ratios were then multiplied with AGC to obtain CRC estimates of all trees in the experiment.

#### Coarse woody debris carbon

All visible branches and stems fallen on the ground were collected annually in each plot and weighted to obtain a measure of coarse woody debris (CWD) biomass. CWD biomass was converted to C (hereafter CWDC) using the species-specific trunk C concentration detailed above.

#### Herbaceous carbon

Herbaceous vegetation was cut, dried, and weighed in four quadrats (0.5 m^2^) per plot (Potvin et al., 2011). C concentration was determined using an elemental analyser, and herbaceous biomass was converted to herbaceous C using the average C concentration (42.72%) of legumes and grasses/non-leguminous herbs.

#### Leaf litter carbon

Leaf litter was collected bi-weekly in 3–6 litter traps of 1 m^2^, with traps positioned 1 m away from a tree of each species present in each plot, see Scherer-Lorenzen et al. (2005). Leaf litter production was calculated by dividing total dry biomass from each trap by the number of days between two litter collection dates to determine the rate of litter fall per day per m^2^. Litter biomass production was converted to litter C production (hereafter ‘litter C’) using plot- and species-specific carbon concentrations from dry season litter (Scherer-Lorenzen et al., 2007).

#### Soil organic carbon

Four soil cores were collected to a depth of 10 cm from each plot during plantation establishment (2001) and in the mid and late periods, dried, and analysed for bulk density, SOC concentration (%), and δ^13^C values. Litter was removed before sampling. SOC (kg m^-2^) stock was calculated from bulk density and C concentration. We examined not only SOC but also its C_3_- and C_4_-derived fractions since the latter is associated with the C_4_ grasses within the herbaceous vegetation and in the pasture that existed prior to the plantation establishment, while the former is associated with the C_3_ inputs via litter of the trees and C_3_ herbaceous plants (Moore et al., 2018). Assuming a C_3_ plant δ^13^C input of -28 ‰ and a residual C_4_ plant δ^13^C of -13 ‰, estimates of the percentage and mass of C_3_-plant derived SOC (SOC_3_) and C_4_-plant derived SOC (SOC_4_) were made (Moore et al., 2018), where SOC_3_ and SOC_4_ are percentages of total SOC that add up to one hundred percent. This approach allowed us to determine the temporal changes in SOC derived predominately from trees (SOC_3_) and C_4_ grasses in the former pasture (SOC_4_).

#### Leaf litter decomposition

Leaf litter decomposition, hereafter litter decomposition, was measured using nylon bags filled with dry litter from litter traps. For species mixtures, equal proportions of litter from each species were used, see Scherer-Lorenzen et al. (2007). Litter decomposition was measured in a subset of five monocultures (one plot for each species), three three-species mixtures, and three five-species mixtures. Mass loss was determined by drying and weighing the remaining litter and the percent mass remaining was recorded.

#### Root decomposition

Root decomposition was measured using the root material of the five tree species using nylon bags filled with dry roots of 4^th^ and 5^th^ orders (Guerrero-Ramírez et al., 2016). Decomposition bags were installed in the ten monocultures and three five-species mixtures using equal proportions of roots from each species in mixtures. Mass loss was determined by washing, drying and weighing the roots and the percent mass remaining was recorded.

#### Soil microbial biomass

C_mic_ was measured as substrate-induced respiration, that is, the respiratory response of microorganisms to glucose addition (Anderson and Domsch, 1978). C_mic_ was calculated according to Beck et al. (1997), see Cesarz et al. (2022).

#### Soil respiration

Total soil respiration (i.e. autotrophic and heterotrophic respiration) was measured with a portable infrared gas analyser equipped with a soil respiration chamber at six to eight randomly chosen locations per plot. Changes in CO_2_ concentration over time were recorded when pressing the chamber gently on the forest floor.

#### Canopy opening

Canopy opening (i.e. canopy gap fraction in %) was measured using four hemispheric photos per plot when trees were fully leaved-out. Photos were analysed by the Gap Light Analyser (GLA) program (Frazer et al., 1999), see Sapijanskas et al. (2014). Canopy opening can be considered a measure of canopy space-filling, which may mediate tree diversity effects on C stocks and fluxes (see Table S1).

### Data Analysis

We expressed the stock and flux variables in a common unit of 1 m^2^ to avoid extrapolating variables that were measured only in small areas. We considered AGC, CRC, SOC and C_mic_ as stock variables. AGC and CRC (kg) of individual trees were summed per plot and then expressed at the scale of one m^2^ to ensure comparability of all measurements and to account for slight variations in plot size. For SOC (initially measured as kg m^-2^), the data were averaged at the plot level for each year. We also calculated changes in stocks between the three observation periods as:

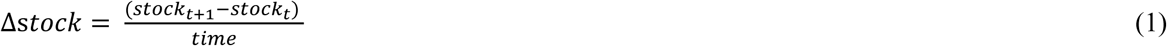

where *stock* is either AGC, CRC or SOC and its fractions SOC_3_ and SOC_4_ in period *t* and *time* is the number of years and months between two measurements (see Fig. S1 for a timeline) resulting in Δstock estimates in kg C m^-2^ year^-1^. We considered CWDC, herbaceousC, litterC, litter decomposition, root decomposition, and soil respiration as annual flux variables. CWDC produced in one year was found to mostly decompose until the end of the wet season in each year of our observation period. Similarly, herbaceousC regrew each year and dead herbaceous material decomposed completely within a year. We therefore considered annual measurements of CWDC and herbaceousC in our tropical forest system as annual fluxes (kg C m^-2^ year^-1^) rather than as stocks, as this attribution more closely reflected the reality in the examined tropical forest compared to calculating changes in these variables between several years. The C flux variables were analysed as follows. CWDC was measured at plot level and down-scaled to 1 m^2^ accounting for plot size. HerbaceousC was averaged at the plot level for each year. LitterC (kg C m^2^ day^-1^) was averaged per plot and across the different collection dates and then scaled to an annual flux (kg C m^2^ year^-1^). The rate of leaf litter and root decomposition (k) per plot and year was calculated based on the percent mass remaining and the days of decomposition (see Supplementary Methods) using a single-pool exponential decomposition model following Adair et al. (2010); decomposition data was not analysed jointly with other variables as decomposition was not measured for all plots. Microbial biomass (µg C_mic_ g^-1^ dry weight soil) and soil respiration (μmol m^-2^ s^-1^) were calculated as average values across measurement locations per plot. Canopy opening was averaged across the different samples within one plot and expressed in %.

#### Multivariate Analyses of Variance

A snapshot of the compartments of the forest C cycle after 16 years of growth (2016–2017) was obtained by Multivariate Analyses of Variance (MANOVA) testing the effect of tree diversity for different compartments expected to be correlated. Two MANOVAs examined the tree biomass related variables (AGC, CRC and CWDC) and the soil related ones (SOC, SOC_4_ and SOC_3_). A third MANOVA analysed the effect of tree diversity on canopy opening and litterC considering that both variables might be correlated. The fourth MANOVA considered soil microbial biomass and soil respiration. The analyses were performed using Proc GLM of SAS version 9.4.

Understanding the build-up of the diversity effects across the examined periods proceeded using mixed-effects Analyses of Variance (mixed-effects ANOVAs). As we were interested in temporal dynamics, we focused on annual C fluxes and changes in stocks rather than on stocks per se to ensure a better comparability of the examined variables and to avoid legacy effects present in stock variables (see e.g. Chen et al. (2023)). The mixed-effects ANOVAs were performed for each Δstock and flux variable at the plot level, according to the following model:

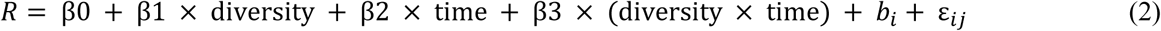

where *R* is the respective response variable and *diversity* had four levels corresponding to the number of species planted per plot (1, 2, 3, and 5), and *time* had three levels (early, mid, and late period). *β0, β1* and *β3* are the fixed effect coefficients, *b_i_* is the random effect for experimental plot accounting for repeated measurements and *ε_ij_* the error term. The model assumes the random effect *b_i_* to be normally distributed with mean and variance of N(0, σ^2^). Our mixed-effects ANOVAs allow testing for the presence of a time by diversity effect, namely a differential build-up of C stocks and fluxes through time in response to tree diversity. ANOVAs were used to test for diversity effects on root decomposition and soil respiration which were only measured once. Extreme values and model assumptions, including normality and heteroscedasticity, were checked visually and with Shapiro-Wilk test (see Supplementary Analysis 1). We decided to only remove two CWDC data points, as in all other cases, a plausible biological explanation existed, and, where necessary, we log-transformed data prior to model fitting to normalize residuals. Mixed-effects ANOVAs were fit in R version 4.3.0 with the packages lme4 (Bates et al., 2015) and lmerTest (Kuznetsova et al., 2017). Least-squares means were estimated with the emmeans package (Lenth, 2020).

#### Structural equation models

To understand the mechanisms underlying tree diversity’s control on C dynamics across time, we used structural equation models (SEMs). We developed a hypothesis-driven conceptual model based on *a priori* knowledge of mechanisms that may drive and relate C stocks and fluxes in forest ecosystems (Fig. S3; Table S1). This approach enabled us to test the direct and indirect relationships between C stocks, fluxes, and tree diversity. Indirect relationships are those that are mediated by other variables. We tested whether tree diversity affected C stocks and fluxes indirectly via diversity-induced decreases in canopy opening due to enhanced canopy space-filling or increases in tree growth through examining relationships between tree species richness, canopy opening, and tree growth (expressed as ΔAGC). We subsequently expected canopy opening and tree growth to influence herbaceousC, litterC and CWDC in that (1) decreased canopy opening at high diversity would correlate negatively with herbaceousC but positively with litterC while (2) enhanced tree growth at high diversity would correlate positively with litterC and CWDC. We did not include direct pathways between tree diversity and these variables as we expected tree diversity effects to be predominately mediated by canopy opening or tree growth. We subsequently expected herbaceousC, litterC, and CWDC to be the main aboveground C inputs to the soil, hypothesizing that they would positively influence and correlate with ΔSOC. Belowground, we included pathways between tree species richness and ΔCRC and between ΔCRC and ΔSOC. As we were interested in tree diversity effects, we focused on SOC_3_, the tree-derived fraction of SOC, but also tested the same SEMs for SOC (sum of SOC_3_ and SOC_4_). Moreover, as we assumed canopy opening and ΔAGC, and ΔCRC and ΔAGC to be correlated, we included partial correlations between these variables. Finally, we tested for potential direct effects of tree diversity on ΔSOC not mediated by the tested relationships. To examine temporal trends, we fit separate SEMs per period. As information on ΔSOC, a crucial variable for our SEMs, was only available for the mid and late periods (see Fig. S1), we fit SEMs only for these two periods.

All SEMs focussed on C fluxes and changes in stocks to ensure a better comparability between variables. Moreover, we only included variables available in kg C m^2^ year^-1^, except for canopy opening. We used piecewise SEMs (Lefcheck, 2016) to test the relative importance of and support for these hypothesized pathways. Global model fit was assessed via Fisher’s C statistic (P>0.05). We assessed the independence of variables and included partial, non-directional correlations to improve model fit based on tests of directed separations (P<0.05 for violation of independence claims). For each SEM we calculated standardized path coefficients, scaled by the standard deviations of the variables, which allowed us to compare the strength of paths within and among models (Lefcheck et al., 2018). Individual pathways were fit as linear models considering the number of species planted per plot (1, 2, 3, and 5) as continuous and not as categorical variable as in the mixed-effects ANOVAs. SEMs were fit with the package piecewiseSEM (Lefcheck, 2016) and linear mixed-effects models with the packages lme4 (Bates et al., 2015) and lmerTest (Kuznetsova et al., 2017) in R version 4.3.0.

## Results

### C stocks and fluxes after 16 years of tree growth: a snapshot in time

Over a period of 16 years, the experimental tree plantation accumulated an average of 35.9±2.7 Mg C ha^-1^ in the trees (AGC + CRC), while SOC decreased on average by 11.2±1.1 Mg C ha^-1^, resulting in a net gain of 24.7±2.9 Mg C ha^-1^ or 90.7±10.6 Mg CO_2eq_ ha^-1^. MANOVA unveiled a significant effect of tree diversity on the C compartments directly related to trees: AGC, CRC and CWDC (Table 1, Fig. 1). The diversity effect was mainly driven by AGC (Fig. 1) with both CRC and CWDC being significantly correlated to AGC (0.708, p=0.0007 and 0.624, p=0.004, respectively). In 2017, according to Tukey Studentized range test, tree AGC in the 5-species plot, was, with 35.7±1.8 Mg C ha^-1^, significantly higher than in monocultures (22.8±3.4 Mg C ha^-1^), a 57% increase (Fig. 1). The MANOVA computed with the canopy-related variables (canopy opening and litterC) also showed a significant effect of diversity (Table 1) with a significant negative correlation between the two variables (-0.741, p=0.0004). This diversity effect was predominantly driven by litterC that was 64% higher in 5-species mixtures than in monocultures. At 1.1–1.4 Mg C ha^-1^ (Fig. 1; Fig. S4) herbaceousC played a role similar to litterC in the system, albeit it did not respond significantly to diversity (Table 2). None of the two MANOVAs performed on soil-related C compartments detected a significant effect of tree diversity (Table 1).

**Fig. 1.**
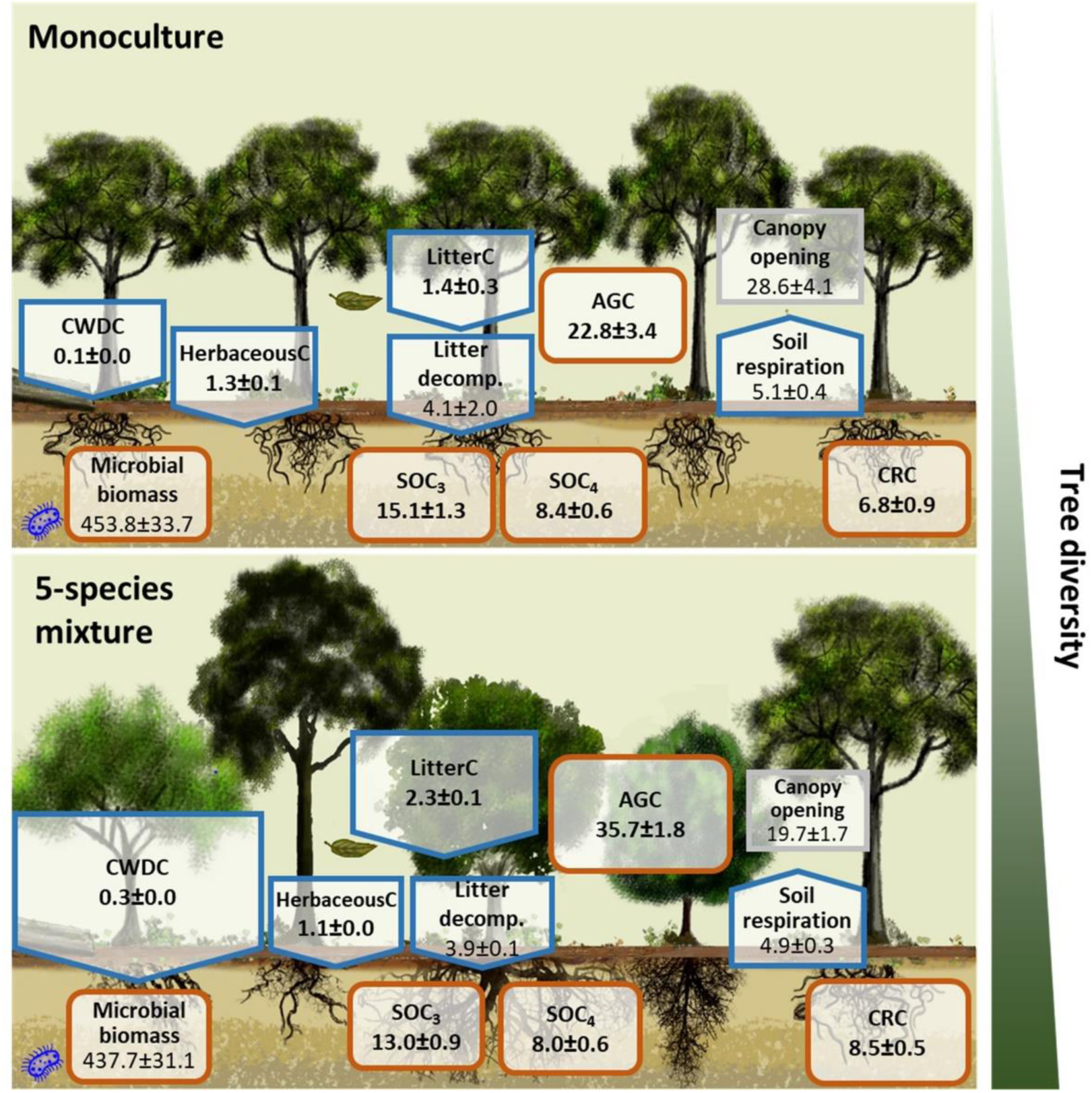
C stocks and fluxes after 16 years of tree growth. Shown are means and standard errors of the C stocks in Mg C ha^-1^ (brown boxes) and fluxes in Mg C ha^-1^ year^-1^ (blue arrows); numbers printed in bold. Variables in other units, including canopy opening in %, litter decomposition rate k year^-1^, soil respiration given in μmol m^-2^ s^-1^ and microbial biomass given in µg C_mic_ g soil dw^-1^ are not printed in bold to allow for separation. The size of the boxes and arrows in the 5-species mixture are scaled relative to the monoculture, with diversity-induced increases or decreases in C stocks and fluxes indicated by larger or smaller boxes/arrows, respectively. An overview of all analysed mixtures (2-, 3- and 5-species mixtures) is shown in Fig. S4. The sum of SOC_3_ and SOC_4_ gives SOC. AGC: aboveground tree C, CRC: coarse root C, CWDC: coarse woody debris C, SOC_4_: C_4_ derived soil organic C (SOC), SOC_3_: C_3_ derived SOC.

**Table 1.** MANOVAs for C stocks and fluxes after 16 years of growth. The main effect tested was tree diversity with 4 different levels (1, 2, 3 and 5 species per plot). Abbreviations are given in Fig. 1. Significant effects (p <0.05) printed in bold.

| Compartments | Roy's Greatest Root | F<sub>3,18</sub> | p-value |
| --- | --- | --- | --- |
| AGC/CRC/CWDC | 0.8975 | 5.39 | **0.0080** |
| LitterC/Canopy opening | 0.5974 | 3.58 | **0.0343** |
| SOC/SOC<sub>4</sub>/SOC<sub>3</sub> | 0.2059 | 1.24 | 0.3260 |
| Microbial biomass/Soil respiration | 0.1722 | 1.03 | 0.4015 |

**Table 2.** Mixed-effects ANOVAs on changes in C stocks and fluxes over time. The analyses considered up to three time-intervals, early, mid, and late period across a total of 16 years, see the timeline in Fig. S1 for details. Abbreviations are as in Figure 1. Entries in the table are F and p values, with p <0.05 *, p< 0.01 **, p<0.001 ***, and ns = not significant. Significant effects printed in bold. Litter decomposition rate was log-transformed prior to model fitting to normalize residuals. For root decomposition rate and soil respiration, ANOVA results are shown.

| C stock/flux | Diversity | Time | Time × Diversity |
| --- | --- | --- | --- |
| ΔAGC | **5.07\*** | **11.25 \*\*\*** | 1.67 ns |
| ΔCRC | 1.11 ns | 2.74 ns | 0.72 ns |
| CWDC | **3.30 \*** | 0.23 ns | 2.01 ns |
| HerbaceousC | 0.98 ns | **47.10 \*\*\*** | 0.37 ns |
| LitterC | 0.81 ns | **100.74 \*\*\*** | **6.25 \*\*\*** |
| ΔSOC<sub>3</sub> | 0.92 ns | **29.10 \*\*\*** | 1.61 ns |
| ΔSOC<sub>4</sub> | 0.67 ns | **44.57 \*\*\*** | **5.35 \*\*** |
| ΔSOC | 0.62 ns | 0.01 ns | 2.55 ns |
| Litter decomposition | 0.20 ns | **4.39 \*** | 1.17 ns |
| Root decomposition | 0.04 ns | - | - |
| Microbial biomass | 0.81 ns | **46.32 \*\*\*** | 0.90 ns |
| Soil respiration | 0.45 ns | - | - |
| Canopy opening | 2.06 ns | **31.73 \*\*\*** | 1.29 ns |

### Diversity effects on changes in C stocks and fluxes across time

With a least-squares mean of 0.140 kg C m^2^ yr^-1^, tree ΔAGC increment was significantly slower in monocultures than in most mixtures, which had increments of 0.140, 0.232, and 0.226 kg C m^2^ yr^-1^ in 2-, 3-, and 5-species mixtures, respectively (Table 2, Fig. 2). The positive tree diversity effect on ΔAGC tended to strengthen over time in the 5-species mixtures (p = 0.16 for the time × diversity interaction). CWDC was significantly lower in monocultures (least-squares mean of 0.010 kg C m^2^ yr^-1^) than in all mixtures, with 0.012, 0.024, and 0.021 kg C m^2^ yr^-1^, for 2-, 3- and 5-species mixtures, respectively. In contrast, ΔCRC increment did not significantly differ between monocultures and mixtures (Table 2, Fig. 2).

**Fig. 2.**
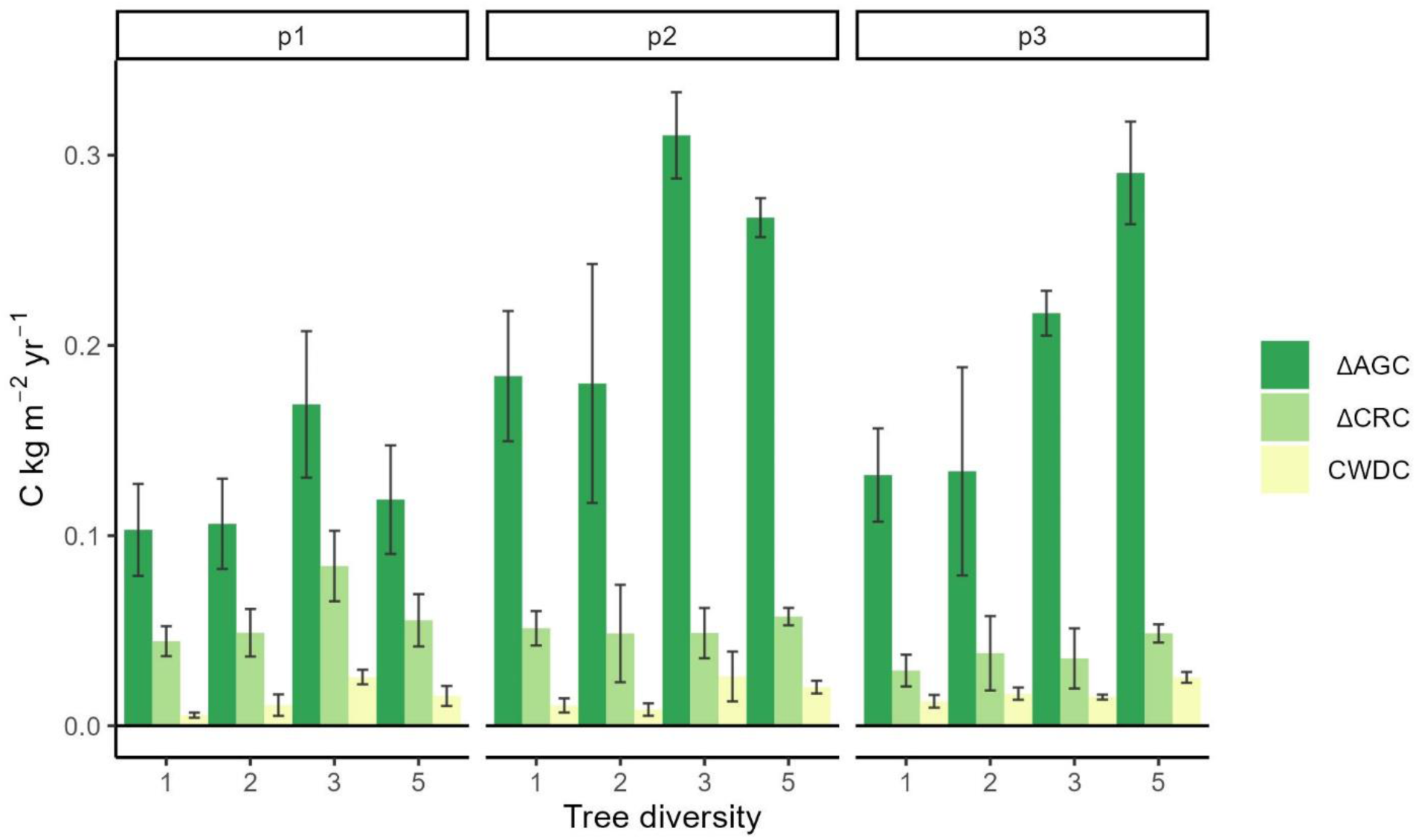
Mean changes in aboveground tree C (ΔAGC), coarse root C (ΔCRC) and coarse woody debris C (CWDC) over time and with diversity. The analyses considered three time-intervals, early (p1), mid (p2), and late period (p3). Coloured bars show means and error bars standard errors of the mean for the examined variables calculated as detailed in Eq. 1 and Fig. S1.

We further compared the strongest aboveground C fluxes to the soil (herbaceousC and litterC, which were an order of magnitude higher than CWDC; Fig. 1) with observed changes in ΔSOC (Fig. 3). Across the plantation, herbaceousC did not vary with diversity but decreased significantly with time, with 0.201, 0.105, and 0.125 kg C m^2^ yr^-1^ for the early, mid, and late period, respectively (Table 2, Fig. 3). Conversely, litterC increased significantly over time, with diversity effects depending on the period of plantation development (Table 2): In the early period, litterC was lowest and similar across diversity levels, while in the mid and late period litterC was lowest in monocultures with 0.149 and 0.141 kg C m^2^ yr^-1^ and highest in 5-species mixtures, with 0.225 and 0.232 kg C m^2^ yr^-1^ (Table 2, Fig. 3).

**Fig. 3.**
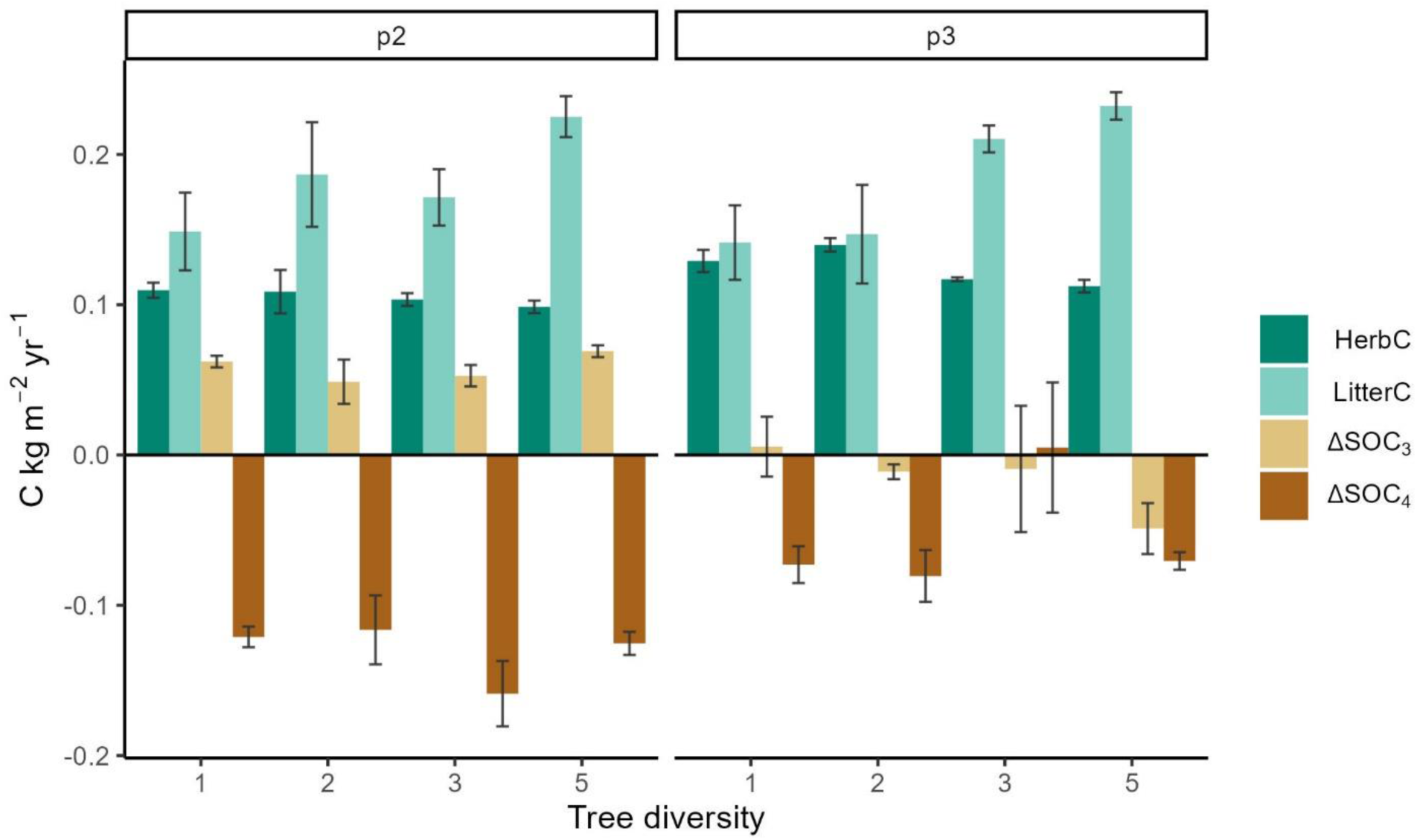
Mean changes of dominant aboveground C fluxes to the soil with time and diversity and corresponding observed changes in soil organic carbon (ΔSOC). Coloured bars show means and error bars standard errors of the mean of herbaceous C and leaf litter C and changes in C_3_ and C_4_-derived SOC (SOC_3_ and SOC_4_). The analyses considered two time-intervals, mid (p2), and late period (p3), as SOC data was only measured in these two periods (Fig. S1). The examined SOC changes were calculated as detailed in Eq. 1 and Fig. S1.

The predominately tree-derived ΔSOC_3_ showed a significant change over time but not with diversity (Table 2): in the mid period, all plots – irrespective of their diversity – showed a net increase in ΔSOC_3_ while most plots did not show any SOC_3_ loss or increment in the late period (Fig. 3). A notable exception are the 5-species mixtures, which tended to lose SOC_3_ in the late period (Fig. 3). The indication of SOC_3_ increment in the mid period coincided with the slowest litter decomposition in this period (Fig. 4). Overall, litter decomposition rates changed significantly over time (being lowest in the mid period) but not with diversity (Table 2; Fig. 4); still, decomposition rates in 5-species mixtures tended to be higher in the late period than in the early and mid period (Fig. 4). High variations in litter decomposition rates in monocultures resulted from the fast decomposition of Hc litter in all periods (Fig. S5). Similarly to litter decomposition, the root decomposition rate did not change with diversity (Table 2, Fig. 4; note that root decomposition was only measured in the mid period).

**Fig. 4.**
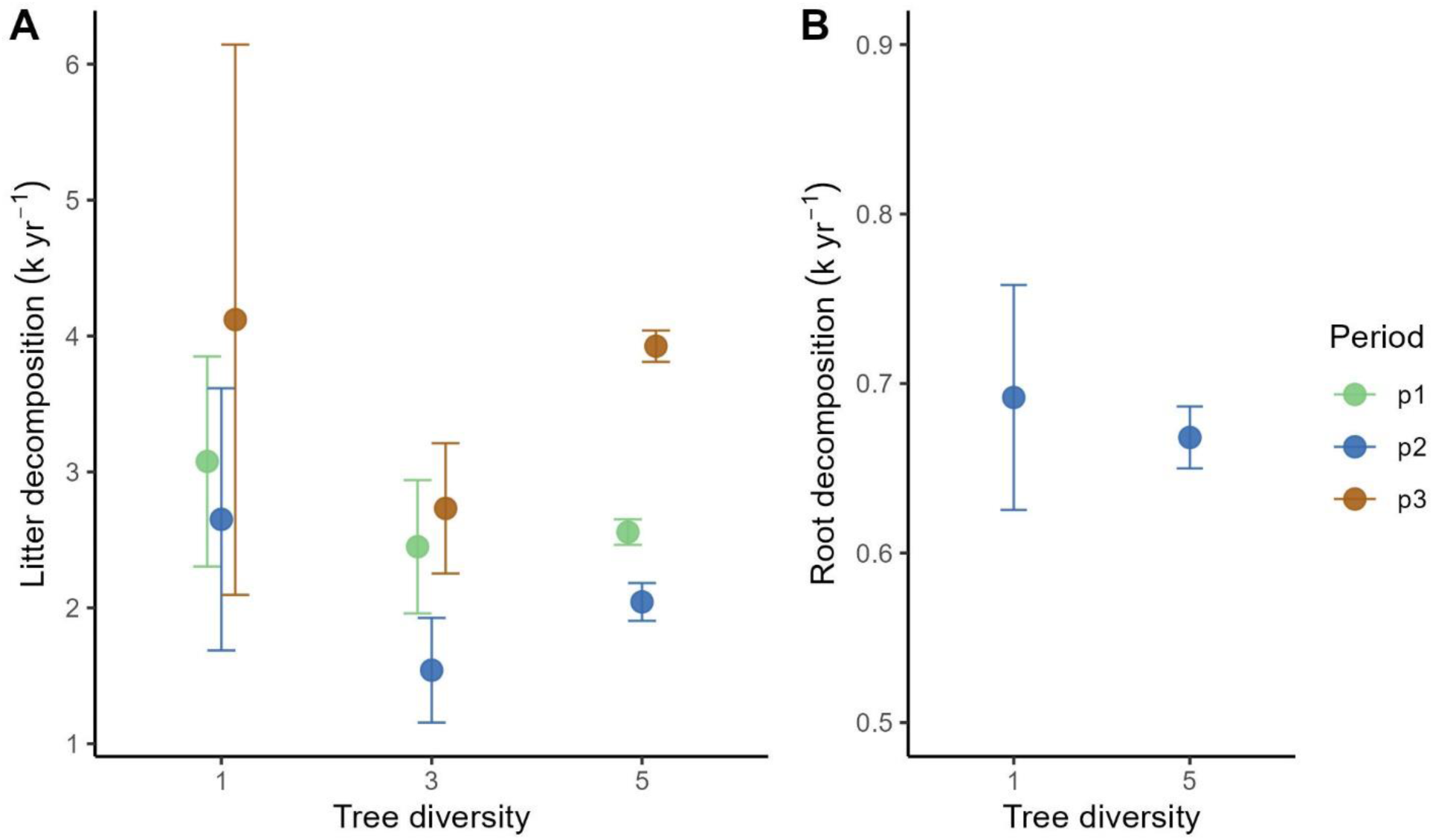
Mean changes in leaf litter and root decomposition rates with time and diversity. Decomposition rates (k yr^-1^) were calculated with a single-pool exponential decomposition model. Coloured points show means and error bars standard errors of the mean. The analyses considered three time-intervals, early (p1), mid (p2), and late period (p3).

ΔSOC_4_, which is associated mainly with the C_4_ grasses that existed before the plantation establishment, varied significantly in response to both time and the time by diversity interaction (Table 2). Between 2001 and 2011 the reduction in C_4_ in the upper 0–10 cm of soil was around twice as fast as between 2011 and 2017 with least-square means of -0.13 kg C m^2^ yr^-1^ and -0.055 kg C m^2^ yr^-1^, respectively (Fig. 3). The significant time by diversity interaction was driven by the three-species mixtures where the decrease in ΔSOC_4_ was strongest in the mid period and disappeared in the late period while the other treatments experienced continued loss in ΔSOC_4_ in the late period (Fig. 3). Overall, reductions in predominately pasture-derived SOC_4_ were larger than any observed gains in tree-derived SOC_3_ resulting in a net negative SOC-balance of reforestation (Fig. 3).

Finally, none of the remaining variables, including microbial biomass, soil respiration and canopy opening responded significantly to diversity (Table 2), even though canopy opening tended to be lower in 5-species mixtures compared to monocultures (F = 2.06, p = 0.14), particularly in the mid- and late period. However, for soil microbial biomass and canopy opening, for which we had repeated measurements, we observed pronounced temporal changes (Table 2). Soil microbial biomass increased significantly from 315 to 453 µg C_mic_ g soil dw^-1^ from the mid to the late period. Not surprisingly, canopy opening, and thus light transmission, declined strongly with progressing stand development from a least-squares mean of 47.8% in the early to 21.1% in the mid period to than increase slightly again to 27.2% in the late period.

### Effects of diversity on C stock-flux relationships

Using structural equation models, we explored the effect of tree diversity on linkages amongst C stocks and fluxes above- and belowground (Fig. 5). In both periods of plantation development (mid and late period), tree diversity significantly decreased canopy opening and increased ΔAGC with standardized path coefficients of -0.41 and 0.42 in the mid period and -0.4 and 0.71 in the late period (Fig. 5). Tree diversity effects on ΔAGC thus increased by ∼70% from the mid to the late period. Canopy opening exerted a significant positive effect on herbaceousC but decreased litterC with standardized path coefficient of 0.57 and -0.40 in the mid period and 0.52 and -0.43 in the late period. ΔAGC significantly increased both litterC and CWDC with standardized path coefficients of 0.51 and 0.68 in the mid period and 0.53 and 0.62 in the late period. Hence, we observed a consistent control of diversity on aboveground C stocks and fluxes and their linkages across time. Aboveground, tree diversity led to a net increase of C fluxes to the soil through its indirect effects on herbaceousC, litterC and CWDC of -0.23, 0.38, 0.29 and -0.21, 0.55, 0.44 respectively for the mid- and late period (see Lefcheck, 2016 for calculation of indirect effects).

**Fig. 5.**
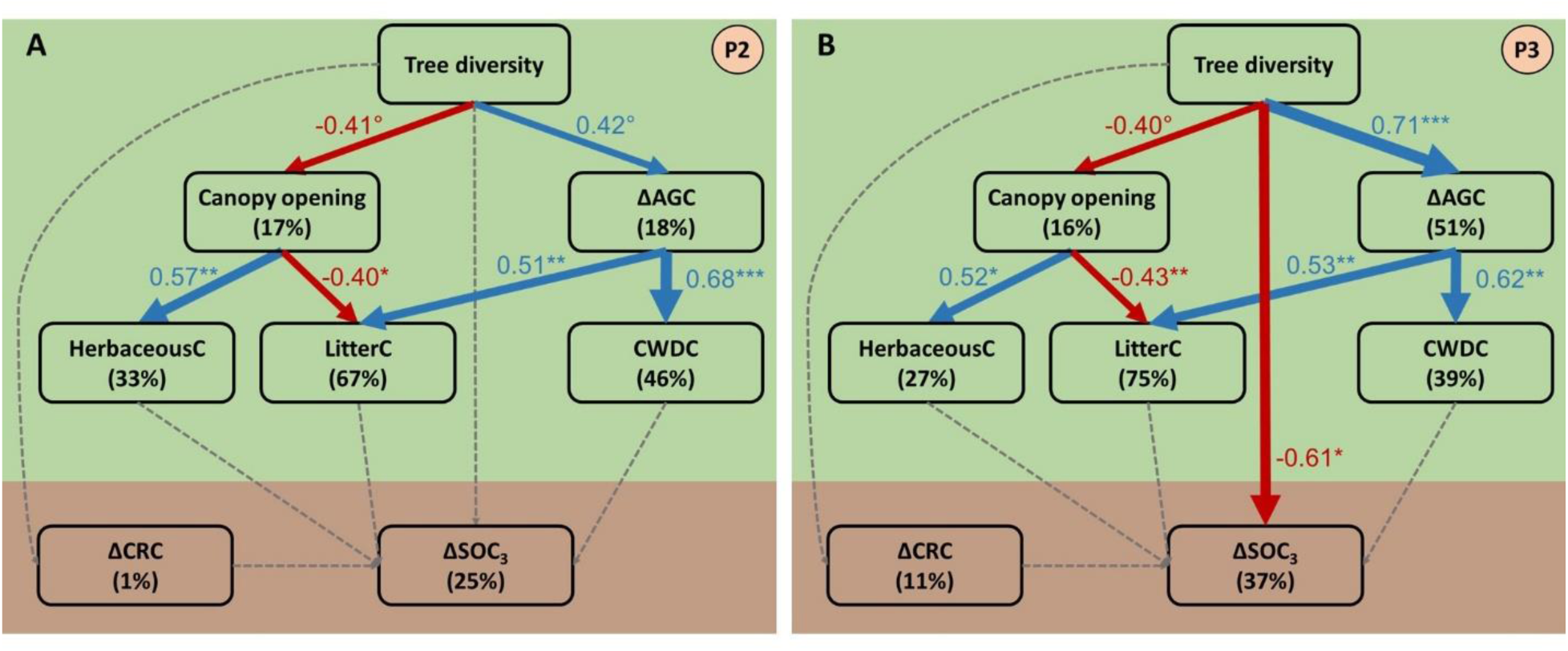
Direct and indirect effects of tree diversity on C stocks and fluxes. The SEMs were fit for the mid period (panel A, P2) and the late period (panel B, P3) and partition potential tree diversity effects on C stocks and fluxes into effects mediated via canopy space filling (expressed as canopy opening) and via aboveground tree C changes (expressed as ΔAGC), which are expected to influence herbaceous C, leaf litter C and coarse woody debris C (CWDC). These latter C fluxes are, in turn, hypothesized to influence changes in predominately tree-derived soil organic C (ΔSOC_3_). All variables were calculated as detailed in Eq. 1 and Fig. S1. The SEMs fit the data well (Fisher’s *C* = 23.09, df = 26, *P* = 0.63, *n* = 22 plots for (A); Fisher’s *C* = 21.70, df = 24, *P* = 0.60, *n* = 22 plots for (B)). Examined variables are shown as boxes and relationships as directional arrows with significant positive effects in blue, significant negative effects in red, and nonsignificant effects in dashed gray. The hypothesis-driven conceptual model is shown in Fig. S3. For each significant relationship, standardized path coefficients are shown next to each path with path-width scaled according to coefficient size and asterisks indicating the significance level (°P < 0.10, *P < 0.05, **P < 0.01, and ***P < 0.001). The variation explained in each variable (*R^2^*) is shown below the variable name. The green and brown font indicates above- and belowground variables.

In contrast, we observed no significant linkages between aboveground and belowground C stocks and fluxes and only one direct effect of diversity on ΔSOC_3_ (Fig. 5). In the mid period we observed no significant effects on ΔSOC_3_ nor ΔCRC. In the late period, the only significant effect was a direct negative effect of diversity on ΔSOC_3_ with a standardized path coefficient of -0.61. ΔCRC did not significantly respond to diversity nor did it significantly influence ΔSOC_3_ in either period. SEMs for ΔSOC instead of ΔSOC_3_ yielded similar results, except for the disappearance of the significant diversity effect in the late period and overall lower explained variation (*R^2^*) in ΔSOC (Fig. S6). Overall, belowground changes in C stocks were largely disconnected from the diversity-controlled C stock and flux network observed aboveground.

## Discussion

As the oldest tropical site of TreeDivNet, the Sardinilla experiment provides a unique opportunity to evaluate temporal changes in C stocks and fluxes in the neotropics. Consistent with our hypothesis (H1), we observed that tree diversity can increase C stocks and fluxes. Specifically, we noted a remarkable increase in AGC stocks driven by tree diversity, with an average 57% increase in AGC in five-species mixtures compared to monocultures after 16 years (35.7±1.8 *vs* 22.8±3.4 Mg C ha^-1^; Fig. 1). This observed effect aligns with a recent meta-analysis demonstrating that mixed-species plots stored more carbon in aboveground biomass than monoculture ones (Warner et al., 2023). Comparing our results with chronosequences of secondary forests in Panama and the wider neotropics, we found that our C stock estimates after 16 years (35.8 and 22.6 Mg C ha^-1^ stored respectively in trees and SOC (0–10 cm)) are comparable. For instance, Neumann-Cosel et al. (2011), working like us in the Panama Canal watershed, reported aboveground ranges of 20.55 Mg C ha^-1^ to 56.45 Mg C ha^-1^ with SOC in the first 0-10 cm adding 27.5±3.1 Mg C ha^-1^. Similarly, Gardon et al. (2020) estimated that actively restored secondary forests in Brazil accumulated approximately 100 Mg biomass ha^-1^ after 16 years, while Rozendaal and Chazdon (2015) reported 104±3.7 Mg biomass ha^-1^ in 10–24-year-old secondary forests in Costa Rica, estimates roughly equivalent to 50 Mg C ha^-1^, and akin to the AGC of our most productive mixture plot. Overall, these findings contribute to our understanding of the build-up of C stocks in tropical forest restoration and emphasize the importance of considering tree diversity in such initiatives.

### Forest stability

The long-term C balance of reforestation in the face of progressing climate change may depend more on forest stability and thus C residence time within the forest than on average C accumulation rates. Here, we understand stability as a forest’s ability to maintain functioning over time despite repeated perturbations, such as climate extremes (Schnabel et al., 2021), which is broadly consistent with the insurance hypothesis (Yachi and Loreau, 1999). In the Sardinilla experiment, observed climate extremes left the ecosystem little time to recover between perturbations. The extreme wet spell of 2010 was followed in 2015 by a severe drought (Fig. S2; see also Browne et al., 2021; Detto et al., 2018), and a hurricane in 2016. As Bhaskar et al. (2018) pointed out, it is important to understand forest resilience in the context of constant climatic disturbances. While our study design does not allow us to disentangle the effects of climate extremes from those of stand development on tree diversity effects, it does allow us to test whether positive tree diversity effects persist or even strengthen over time despite repeated climate extremes. Consistent with our hypothesis (H2), and despite these repeated extremes, we uncovered significant positive effects of tree diversity on all the aboveground C stocks and fluxes (Fig. 5). Remarkably, the positive tree diversity effect not only persisted but tended to strengthen over time, at least for ΔAGC. Our previous findings (Hutchison et al., 2018; Schnabel et al., 2019) indicated that tree diversity increased the stability of tree productivity by enhancing growth, buffering temporal variations in growth, and reducing mortality vis-à-vis monocultures. Although two- and three-species mixtures were the most productive in the early period of the Sardinilla experiment (e.g. Healy et al., 2008; Scherer-Lorenzen et al., 2007), five-species mixtures outperformed the less diverse mixtures in later years, as already observed in previous reports (Guillemot et al., 2020; Schnabel et al., 2019). Here, we moved beyond these earlier studies and similar findings in other ecosystems (e.g. Jucker et al., 2014; Schnabel et al., 2021), which focussed on single ecosystem functions, and leveraged a unique dataset on multiple C stocks and fluxes and their interrelationships. This approach allowed us to adopt an integrated ecosystem perspective on C stability in mixed compared to monospecific planted forests. C stability is particularly important considering that tree diversity increased C stocks and fluxes only aboveground, where C is particularly susceptible to climate-driven forest disturbances.

### Relationships between carbon stocks and fluxes

Using structural equation modelling we showed that the tree diversity directly or indirectly effected all the aboveground C stocks and fluxes that we measured (Fig. 5), suggesting that the aboveground components of the C cycle are linked. Canopy opening, a proxy for canopy space filling, played a central role in our SEMs with denser foliage at high diversity enhancing litterC but reducing herbaceousC. Tree architecture apparently plays a crucial role in explaining this effect. After 16 years of growth, trees growing in mixture allocated a higher proportion of their biomass to branches compared to the same species growing in monocultures (Guillemot et al., 2020), which is consistent with higher canopy space filling reported in other tree diversity experiments (Kunz et al., 2019; Williams et al., 2017). Similarly, tree productivity (captured here as ΔAGC) increased with tree diversity, as has been previously reported from our (Guillemot et al., 2020; Schnabel et al., 2019) and other tree diversity experiments (Guerrero-Ramírez et al., 2017). Enhanced tree productivity at high diversity, in turn, enhanced both litterC and CWDC and thus C fluxes to the soil. Likely drivers of these observed positive diversity effects aboveground are complementary species interactions in mixtures, such as higher community-level light capture or complementary water and nutrient uptake from different soil layers reported in Sardinilla (Oelmann et al., 2010; Sapijanskas et al., 2014; Schwendenmann et al., 2015; Zeugin et al., 2010). These positive diversity effects may be particularly pronounced in mixtures of the Sardinilla experiment that feature species with distinctly different growth rates and shade tolerances, which should promote crown complementarity and thus light capture and use efficiency (Forrester, 2017; Potvin et al., 2011; Schnabel et al., 2019). Litter manipulation experiments in tropical forests support the idea of significant leaf litter-driven carbon and nutrient fluxes to the soil (Cusack et al., 2018; Sayer et al., 2024; Wood et al., 2009). After ten years of manipulation, soil C was significantly higher in the 0–5 cm layer in litter addition plots (Cusack et al., 2018). However, litter addition did not strongly impact tree growth (Sayer et al., 2024; Wood et al., 2009). The authors explained their results by the fast carbon cycling and high C input into the soil in tropical forests. Our SEMs did not confirm such positive link between litterC and the predominately tree-derived fraction of SOC (SOC_3_) nor further linkages of SOC_3_ with herbaceousC, CWDC or CRC (Fig. 5). However, we observed a direct negative effect of diversity on SOC_3_ in the late period in which 5-species mixtures tended to loose SOC_3_ (Fig. 3), which may be related to the comparably fast litter decomposition in this treatment during that period (Fig. 4). Overall, this means that the surplus of C fluxes to the soil in the high-diversity mixtures was largely not incorporated into the soil matrix, and thus, the multiple direct and indirect relationships of aboveground C stocks and fluxes largely did not extend belowground.

### Soil organic carbon

Overall, the Sardinilla planted forest gained significant C aboveground, but we observed a loss of SOC in the 0–10 cm layer associated with a reduction in both bulk density and SOC concentration (Table S2). This loss occurred regardless of tree species richness levels, indicating that tree diversity may only sometimes exert a significant effect on SOC, as noted also in another temperate tree diversity experiment (Martin-Guay et al., 2022). Because the planted forest was established on an active pasture (Scherer-Lorenzen et al., 2005), we suggest that the removal of cows likely minimized compaction which resulted in decreased bulk density (Blanco Sepúlveda and Nieuwenhuyse, 2011). Several studies have identified the misrepresentation of changes in SOC mass associated with sampling to a fixed depth where there have been significant changes in bulk density to that depth (e.g. von Haden et al., 2020). Although variable among the treatments and the two sampling periods, comparison of relative changes in SOC concentration and bulk density suggest that about half of the apparent loss in SOC mass is associated with a loss of SOC concentration and the other half with a decrease in bulk density (Table S2) and thus the loss of SOC mass in the plantation may be an overestimate (Fig. 3). However, SOC concentrations in the 10 – 50 cm depth in the pasture prior to the establishment of the Sardinilla experiment ranged from 1 to 2% and a δ^13^C value between -17 and -21‰, suggesting a strong proportion of SOC_4_, and thus there may be further losses in SOC from the subsoil (Moore et al., 2018), as also suggested by (Quartucci et al., 2023). The literature fails to reach a consensus about how reforestation affects SOC (Laganière et al., 2010), with climatic zone, species planted, clay content, past land use, and soil parent material all affecting the SOC balance of forest regrowth (Araujo et al., 2017; Wallwork et al., 2022). For example, in a nearby Panamanian site, SOC stocks did not vary along a chronosequence of secondary forests (Neumann-Cosel et al., 2011) and in Brazil’s Atlantic forest, SOC stock (0–10 cm) of 5-year old re-growing forests was half that of the remnant forest and similar to that of pastures (Zanini et al., 2021). Marín-Spiotta et al. (2009), who reported no changes in SOC during secondary forest establishment in Puerto Rico, explained this by a loss of pasture-derived SOC_4_ which counterbalanced tree-derived gains in SOC_3_. In a meta-analysis, Don et al. (2011) highlighted substantial SOC gain (on average +17.5%) 28 years after land-use change from grassland to secondary forest in the tropics. It is therefore possible that within the next decade a link between tree diversity and SOC emerges in the Sardinilla planted forest, which highlights the need for long-term studies in tree diversity experiments. We propose that Sardinilla’s clay-rich Cambisols, Tropudalfs and Vertisols (average clay content reaches 65% (Moore et al., 2018)) might have amplified compaction related to grazing while the establishment of trees likely loosened up the soil (Table S2) with follow-up effects like an increase in mineralization and thus, increased C loss. The land-use change effect may thus have (partly) overruled the effects of tree diversity and its C_3_-derived C inputs to SOC stock. Given that high soil clay content is common in the tropics (http://hydro.iis.u-tokyo.ac.jp/~sujan/research/gswp3/soil-texture-map.html; Rasmussen et al., 2018) including soil C pool in the C assessments of forest restoration is likely to improve overall C storage estimates (Quartucci et al., 2023).

### Relevance for forest restoration

The importance of forest restoration for ecosystem C storage, a key ecosystem service for climate regulation, has not only been discussed in the scientific literature but has also triggered international commitment to reforestation. As we work on understanding how forest C stocks are being rebuilt through time at the ecosystem level, it is important to remain realistic about the potential of forest restoration to contribute towards mitigating climate change. Sequestering C is a slow process (Baldocchi and Penuelas, 2019): the average yearly net CO_2_ uptake in Sardinilla was 5.67 Mg CO_2eq_ ha^-1^ yr^-1^ or 1.54 Mg C ha^-1^ yr^-1^. To illustrate, we estimated the emissions from a single one-way flight between Frankfurt and Panama City to 62.7 Mg CO_2eq_ using the [ICAO](https://www.icao.int/ENVIRONMENTAL-PROTECTION/CarbonOffset/Pages/default.aspx) carbon calculator. Hence, this flight demands a flux equivalent to that sequestered by ∼11 ha in the Sardinilla planted forest in one year. Some have suggested that the enthusiasm for nature-based solutions risks putting excessive pressure on land use. For example, Dooley et al. (2022) estimated that countries’ climate pledges for land-based carbon dioxide removal would demand 1.2 billion ha of land, an area globally equal to that used to grow food. Our thorough ecosystem-level analysis of C stocks and fluxes sheds some light on the challenges of using active reforestation projects to compensate for emissions. We note that Griscom et al. (2017) used higher sequestration potentials of 2.8 to 4.7 Mg C ha^-1^ yr^-1^ to estimate the potential of reforestation to act as a nature-based solution for climate mitigation. Therefore, while nature-based solutions are undeniably important for C sequestration and other ecosystem benefits, such as biodiversity and water regulation (Griscom et al., 2017; Seddon, 2022), co-benefits and potential trade-offs should be carefully assessed (Schuldt et al., 2023). Mixed planted forests as a nature-based solution may not only enhance carbon stocks and fluxes vis-à-vis monocultures, as we show here, but also decrease the restored forests susceptibility to stress and disturbances and, thereby, increase C permanence (Anderegg et al., 2020) while also providing higher levels of biodiversity and a broader range of ecosystem services (Messier et al., 2021).

## Supporting information

Supplementary Material for Schnabel et al. 2024

## Acknowledgements

We thank all fieldworkers involved in the Sardinilla experiment over the past 16 years and Daniel Lesieur for data management. For soil analyses, field and laboratory support was provided by Muriel Abraham and Meaghan Murphy. FS acknowledges support by the International Research Training Group TreeDì funded by the Deutsche Forschungsgemeinschaft (DFG, grant 319936945/GRK2324). JG acknowledges support by the French Agricultural Research Centre for International Development (Cirad, CRESI program). NE acknowledges funding by the DFG (German Centre for Integrative Biodiversity Research, FZT118; and Gottfried Wilhelm Leibniz Prize, Ei 862/29-1). TM was supported by the Natural Sciences and Engineering Research Council of Canada (NSERC) and the FQRNT-supported Centre for Climate and Global Change Research at McGill University. NRGR and ITH acknowledge support from Fonds de recherche du Québec nature et technologies, NSERC, Quebec Centre for Biodiversity Science, and the DFG (grant 316045089/GRK 2300). CP acknowledges support from NSERC, the Canada Research Chair Program, and the Smithsonian Tropical Research Institute.

## Author contributions

CP and FS conceived the goals and aims of the study and coordinate the Sardinilla experiment. FS, CP, JG, KEB, SC, NE, and CW developed the methodology. CP, FS, JG, MB, SC, NE, TG, NRGR, ITH, CM, LM, JM, TM, YO, MSL, LS, and AW collected and curated the data. FS and CP analysed the data. FS created the figures. FS and CP wrote the original manuscript draft and all authors contributed to reviewing and editing the manuscript.

## Conflict of interest statement

The authors declare no conflict of interest.

## Data availability

The data that support the findings of this study are openly available in Dryad at http://doi.org/tbd. Data will be made openly available upon acceptance.

## Notes

### Competing Interest Statement

The authors have declared no competing interest.

### Summary of Updates

This is revised version 1 of the manuscript, including among others updated SEMs and more comprehensive data on decomposition. The manuscript figures and text have been updated accordingly.

