## Supplementary Material for Schnabel et al. 2024 for "Tree diversity increases carbon stocks and fluxes above- but not belowground in a tropical forest experiment"

**This file includes:**

Figs. S1 to S6

Tables S1 to S3

Supplementary Methods

Supplementary Analysis

Supplementary References

### Measurement periods of C stocks and fluxes

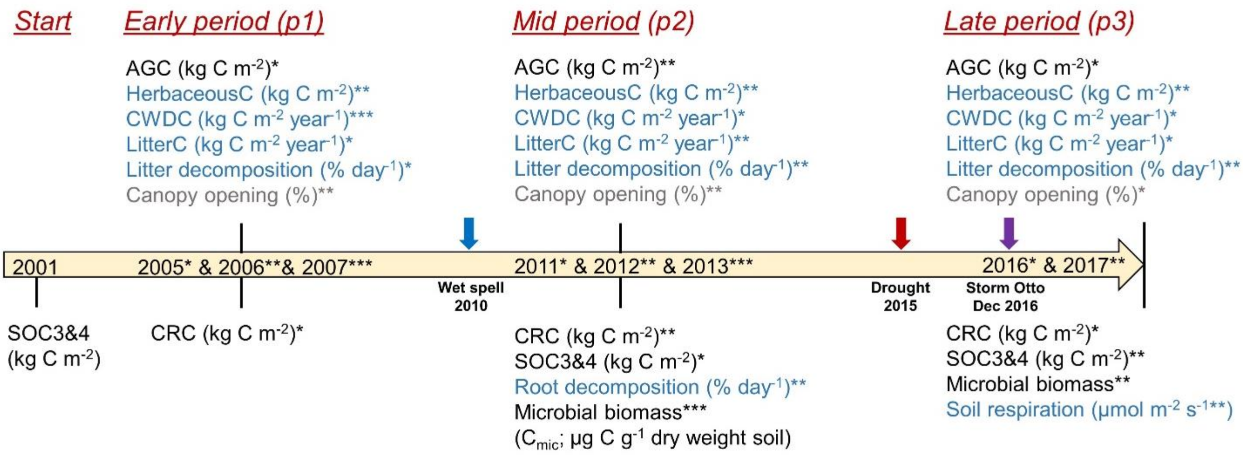

Fig. S1 Data overview. The figure shows the available C stock (in black) and C flux (in blue) measurements for each of the examined periods (early-, mid- and late period). Units for each C stock or flux are given behind the variable's name. The year of each C flux measurement in each period is indicated by the respective asterisk ('*', '**', or '***'). For C stock variables, ΔC stocks were calculated as changes between periods (see Eq. 3) with ΔAGC and ΔCRC expressed as change from 2001–2005 (early period), 2005–2012 (mid period) and 2012–2016 (late period), $\Delta SOC/\Delta SOC_3/\Delta SOC_4$ expressed as change from 2001–2011 (mid period) and 2011–2017 (late period). The mid period was characterized by a an extremely wet event known as *La Purissima* in December 2010 and the late period by a severe drought in 2015 and tropical storm Otto in December 2016, the first Hurricane to ever hit Panama (see Fig. S2 for climate information). AGC: aboveground tree C, CRC: tree coarse root C, CWDC: coarse woody debris C, herbaceousC: C in herbaceous biomass, litter: leaf litter C production, $SOC_4$: $C_4$ derived soil organic C (SOC), $SOC_3$: $C_3$ derived SOC, litter decomposition: leaf litter decomposition, $C_{mic}$: soil microbial biomass C.

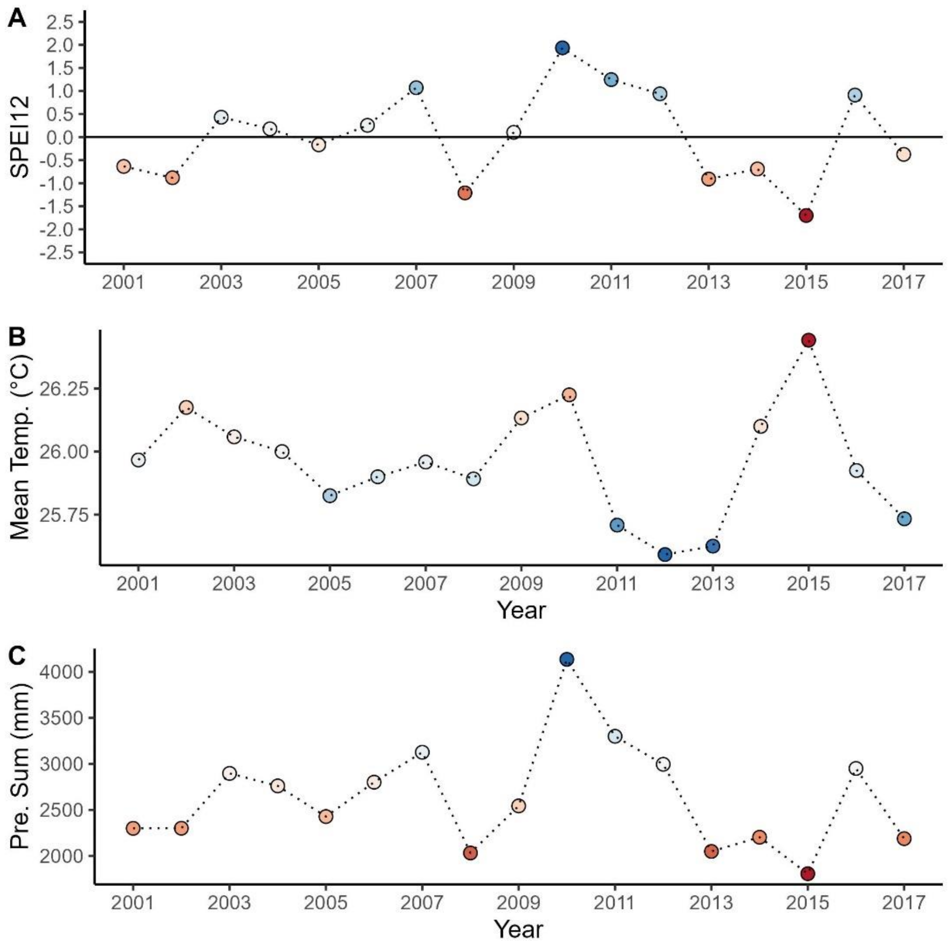

Fig. S2 Standardized Precipitation Evapotranspiration Index (SPEI; Vicente-Serrano et al., 2010) (A), mean monthly temperature (B) and annual precipitation sum (C) per year from 2001–2017. Points are coloured according to their value with deeper red indicating increasing drought severity. Shown are SPEI12 values, i.e. SPEI values calculated for 12 months from January–December. The horizontal line in (A) shows the mean during the observation period (2001–2017), with negative values indicating water deficits and positive values water surpluses. SPEI12 values below -1 and above 1 can be considered as exceptionally dry and wet, respectively (McKee et al., 1993). SPEI was calculated with the SPEI package (Beguería and Vicente-Serrano, 2017) in R from monthly precipitation (mm) and potential evapotranspiration (mm) data. Climate data were obtained from Barro Colorado Island (BCI), the climate station located closest to the Sardinilla experiment (Meteorological Data - Steven Paton, Physical Monitoring Program, Smithsonian Tropical Research Institute, Pers. Com. and ACP Data - Meteorology & Hydrology Dept., Panama Canal Authority).

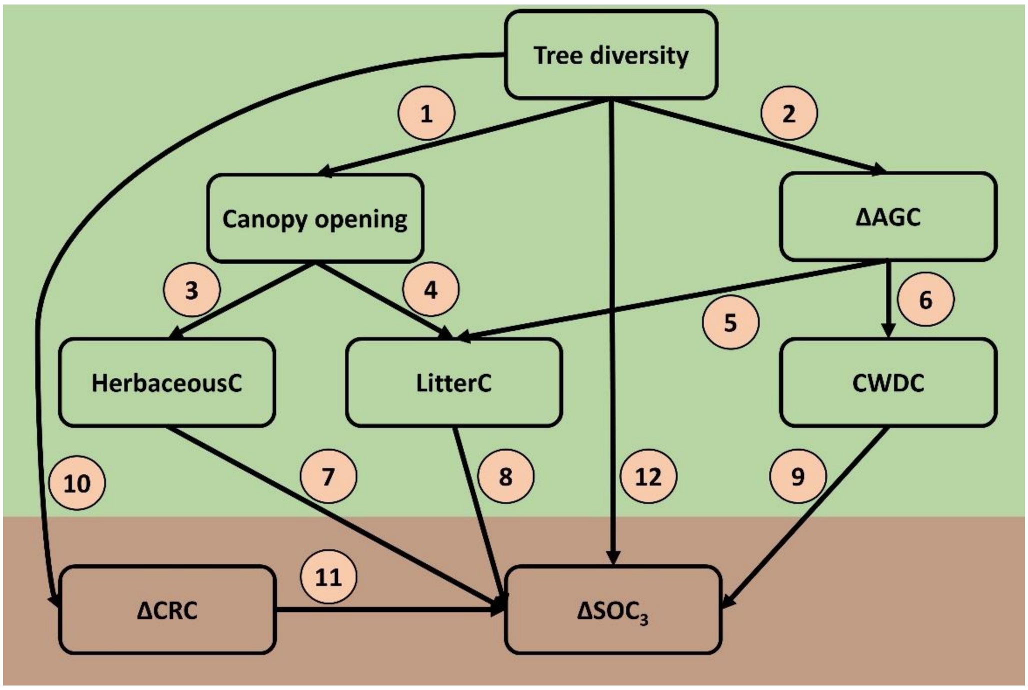

Fig. S3 Structural equation model (SEM) framework. See Supplementary Table S1 for hypotheses on individual relationships (paths 1–12). AGC: aboveground tree C, CRC: tree coarse root C, CWDC: coarse woody debris C, herbaceousC: C in herbaceous biomass, litterC: leaf litter C production, $SOC_3$: $C_3$ derived SOC.

Table S1 Hypothesized relationships between the considered C stocks and fluxes.

| Path | Pathway | Potential mechanism |
|---|---|---|
| **Aboveground** | | |
| 1 | Diversity → Canopy opening | Species richness increases canopy space filling (Jucker et al., 2015; Kunz et al., 2019; Williams et al., 2017) thereby reducing canopy opening. |
| 2 | Diversity → ΔAGC | Species richness increases aboveground tree productivity assessed here as ΔAGC (Guerrero-Ramírez et al., 2017; Huang et al., 2018; Jucker et al., 2020). |
| 3 | Canopy opening → HerbaceousC | Herbaceous biomass increases with increasing canopy opening (Germany et al., 2021). |
| 4 | Canopy opening → LitterC | Leaf litter production decreases with increasing canopy opening (Lin et al., 2015). |
| 5 | ΔAGC → LitterC | Increasing tree productivity increases leaf litter production (Beugnon et al., 2023; Liu et al., 2018; Sapijanskas et al., 2013). |
| 6 | ΔAGC → CWDC | Increasing tree productivity increases coarse woody debris production (Liu et al., 2018). |
| **Belowground** | | |
| 7 | HerbaceousC → ΔSOC | C input via herbaceousC increases SOC (Lange et al., 2015; Weisser et al., 2017), in our study system particularly its $C_4$ fraction ($SOC_4$). |
| 8 | LitterC → ΔSOC | C input via tree litterC increases the $C_3$ fraction of SOC ($SOC_3$) (Castellano et al., 2015; Craig et al., 2022; Cusack et al., 2018; Kuzyakov and Domanski, 2000). |
| 9 | CWDC → ΔSOC | C input via CWDC increases the $C_3$ fraction of SOC ($SOC_3$) (Lodge et al., 2016; Magnússon et al., 2016). |
| 10 | Diversity → ΔCRC | Tree diversity increases CRC though enhanced root biomass production or root carbon concentration (Chen et al., 2018; Eisenhauer et al., 2017; Fornara and Tilman, 2008; Schuster et al., 2023) |
| 11 | ΔCRC → ΔSOC | C input via root carbon residues, which likely increases with CRC, increases the $C_3$ fraction of SOC ($SOC_3$) (Lange et al., 2015). |
| 12 | Diversity → ΔSOC | Tree diversity may influence SOC via mechanisms not considered here. |

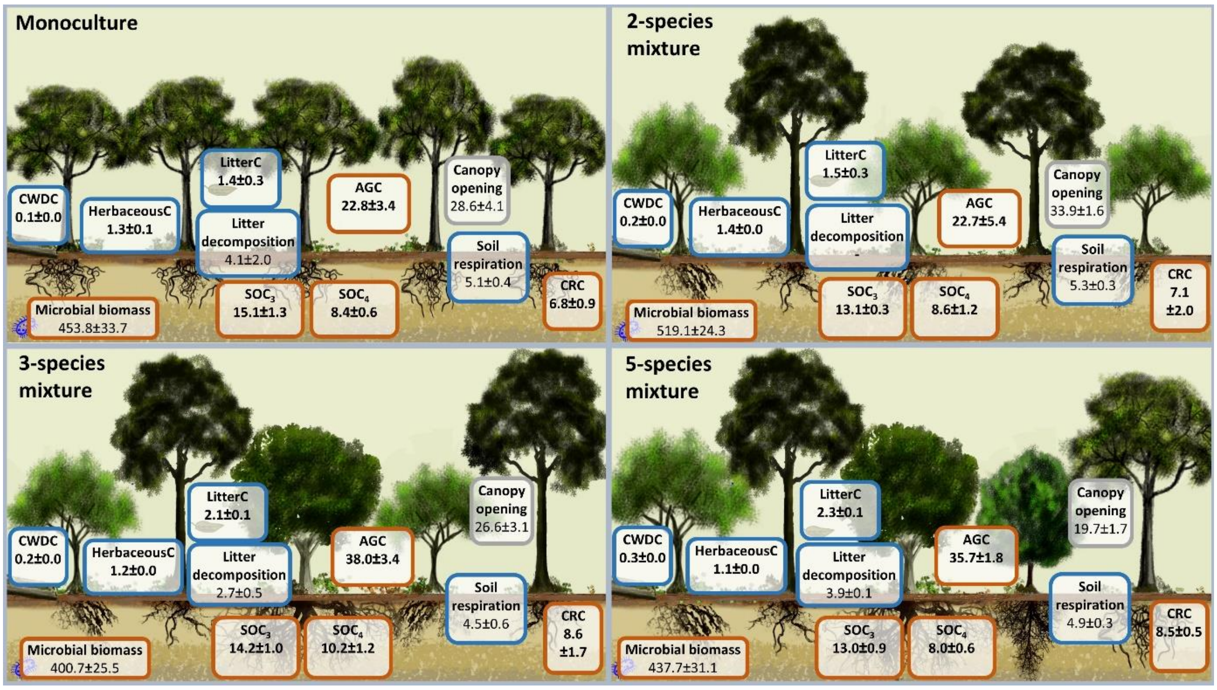

Fig. S4 C stocks and fluxes after 16 years of tree growth. Shown are means and standard errors of the C stocks in Mg C $ha^{-1}$ (brown boxes) and fluxes in Mg C $ha^{-1}$ $year^{-1}$ (blue boxes); numbers printed in bold. Variables in other units, including canopy opening in %, litter decomposition rate k $year^{-1}$, soil respiration given in µmol $m^{-2}$ $s^{-1}$ and microbial biomass given in µg $C_{mic}$ g soil $dw^{-1}$ are not printed in bold to allow for separation. The sum of $SOC_3$ and $SOC_4$ gives SOC. AGC: aboveground tree C, CRC: coarse root C, CWDC: coarse woody debris C, $SOC_4$: $C_4$ derived soil organic C (SOC), $SOC_3$: $C_3$ derived SOC.

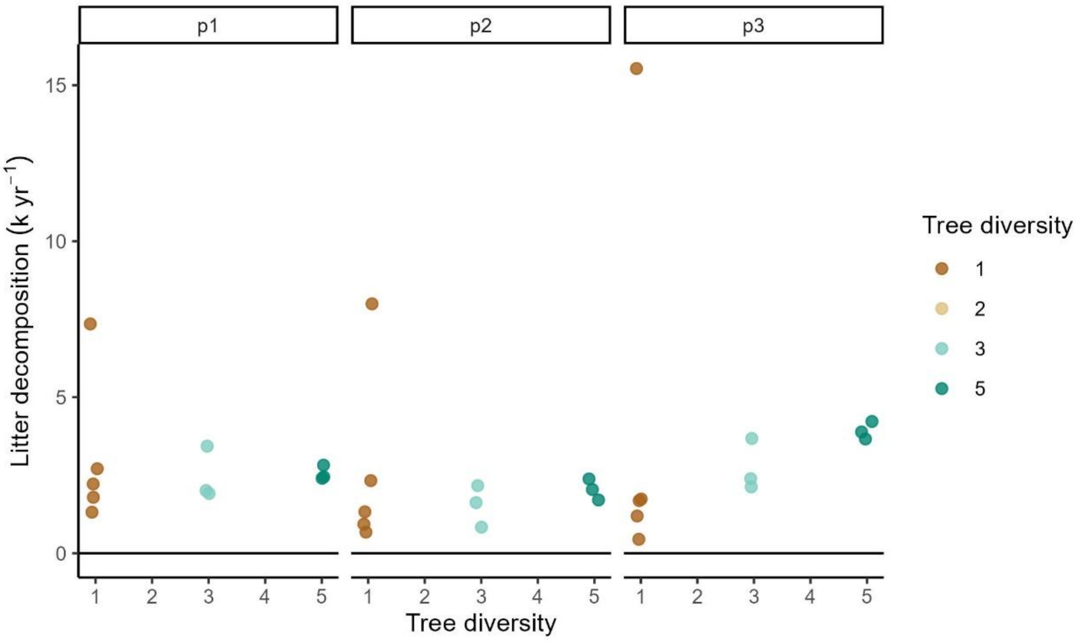

Fig. S5 Leaf litter decomposition rates per plot with time and diversity. Decomposition rates (k year$^{-1}$) were calculated with a single-pool exponential decomposition model. Coloured points show the data for each plot and where coloured according to their diversity level. The analyses considered three time-intervals, early (p1), mid (p2), and late period (p3). The highest decomposition rates in each period correspond to the monoculture of *Hura crepitans* L. (Hc; topmost brown point in each period), featuring easily decomposable litter and far higher decomposition rates than the other species, which might be attributable to its comparably low fiber content as reported by Scherer-Lorenzen et al. (2007).

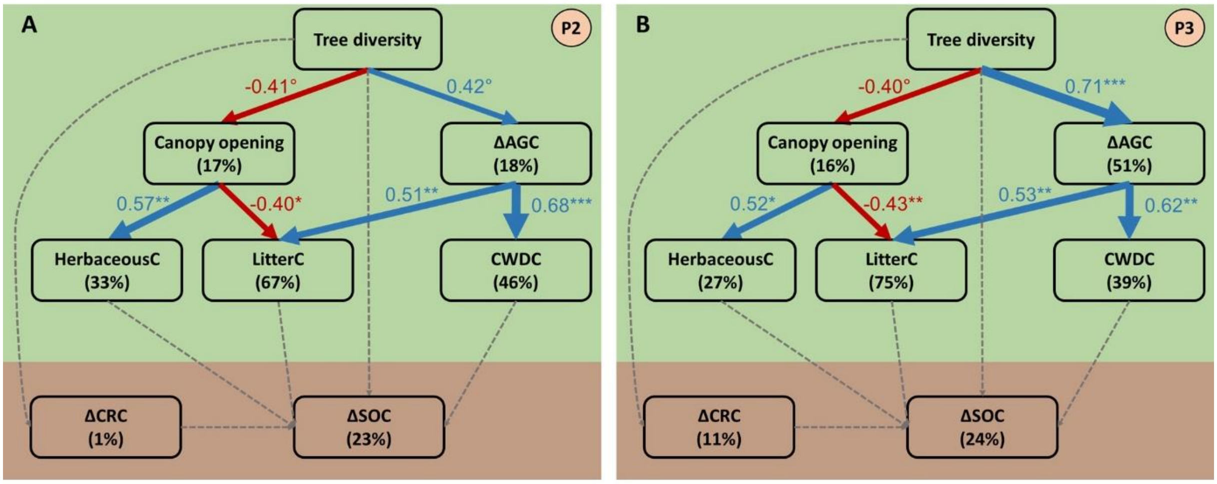

Fig. S6 Direct and indirect effects of tree diversity on C stocks and fluxes considering SOC instead of $SOC_3$ (Fig. 5). The SEMs partition potential tree diversity effects on C stocks and fluxes into effects mediated via canopy space filling (expressed as canopy opening) and via aboveground tree C changes (expressed as ΔAGC), which are expected to influence herbaceous C, leaf litter C and coarse woody debris C (CWDC). These latter C fluxes are, in turn, hypothesized to influence changes in soil organic C (ΔSOC). SEMs were fit for the mid period (panel A, P2) and the late period (panel B, P3) with all variables being calculated as detailed in Eq. 3 and Fig. S1. The SEMs fit the data well (Fisher's $C$ = 20.04, df = 26, $P$ = 0.79, $n$ = 22 plots for (A); Fisher's $C$ = 23.99, df = 24, $P$ = 0.46, $n$ = 22 plots for (B)). Examined variables are shown as boxes and relationships as directional arrows with significant positive effects in blue, significant negative effects in red, and nonsignificant effects in dashed gray. The hypothesis-driven conceptual model is shown in Fig. S3. For each significant relationship, standardized path coefficients are shown next to each path with path-width scaled according to coefficient size and asterisks indicating the significance level (°P < 0.10, *P < 0.05, **P < 0.01, and ***P < 0.001). The variation explained in each variable ($R^2$) is shown below the variable name. The green and brown font indicates above- and belowground variables.

Table S2. Soil characteristics (0 to 10 cm) of the Sardinilla plantation plots as overall mean based on 4 samples collected from within each plot, the number of plots sampled indicated, and standard deviation in parentheses. Treatments are for five mono-specific plots (Ae, Co, Hc, Ls and Tr) and 2-species, 3-species and 5-species plots and sampling was made in 2001, 2011 and 2017. Changes in SOC concentration and bulk density are expressed as a percentage from 2001 to 2011 and from 2011 to 2017. The proportion of SOC mass change derived from SOC concentration is based on partitioning the percentage changes of SOC concentration and bulk density over these two periods. The soil $\delta^{13}C$ values are converted into an estimate of the $C_3$-derived organic matter, based on a $C_3$ plant $\delta^{13}C$ input of -28 ‰ and a residual $C_4$ plant $\delta^{13}C$ of -13 ‰.

| Treatment (# plots sampled) | Year | SOC concentration (mg $g^{-1}$) | Dry bulk density (g $cm^{-3}$) | SOC mass (kg $m^{-2}$) | Change in SOC concentration (%) | Change in bulk density (%) | Proportion of SOC mass change derived from SOC % change | $\delta^{13}C$ (‰) | $SOC_3$ % |
|---|---|---|---|---|---|---|---|---|---|
| Ae (2) | 2001 | 69.4 (19.5) | 0.54 (0.06) | 3.69 (0.70) | | | | -16.8 (1.7) | 25.3 (11.6) |
| | 2011 | 57.2 (9.0) | 0.52 (0.07) | 2.94 (0.51) | -18 | -4 | 0.83 | -21.2 (0.7) | 54.4 (4.5) |
| | 2017 | 52.0 (8.1) | 0.49 (0.05) | 2.58 (0.62) | -9 | -6 | 0.61 | -23.5 (1.6) | 70.2 (10.6) |
| Co (2) | 2001 | 46.6 (7.0) | 0.71 (0.10) | 3.23 (0.28) | | | | -17.1 (1.1) | 27.5 (7.1) |
| | 2011 | 45.0 (6.4) | 0.59 (0.08) | 2.61 (0.32) | -3 | -17 | 0.17 | -21.3 (0.9) | 55.1 (6.4) |
| | 2017 | 43.5 (12.6) | 0.49 (0.04) | 2.12 (0.64) | -3 | -17 | 0.16 | -22.9 (1.4) | 65.9 (9.1) |
| He (2) | 2001 | 59.8 (11.2) | 0.46 (0.07) | 3.34 (0.60) | | | | -16.9 (0.9) | 25.9 (6.0) |
| | 2011 | 53.4 (6.2) | 0.51 (0.04) | 2.71 (0.22) | -11 | 11 | 0 | -21.6 (1.5) | 57.0 (10.2) |
| | 2017 | 44.8 (6.5) | 0.44 (0.06) | 1.95 (0.33) | -16 | -14 | 0.54 | -22.2 (1.1) | 61.3 (7.1) |
| Ls (2) | 2001 | 57.1 (13.5) | 0.57 (0.09) | 3.23 (0.71) | | | | -16.6 (0.6) | 24.0 (3.8) |
| | 2011 | 50.4 (6.2) | 0.54 (0.04) | 2.71 (0.28) | -12 | -5 | 0.69 | -20.3 (1.2) | 48.5 (8.0) |
| | 2017 | 49.5 (3.9) | 0.49 (0.065 | 2.45 (0.46) | -2 | -9 | 0.16 | -22.0 (0.5) | 59.9 (3.4) |
| Tr (2) | 2001 | 56.9 (12.7) | 0.60 (0.08) | 3.40 (0.75) | | | | -16.9 (1.1) | 25.9 (7.3) |
| | 2011 | 49.6 (9.6) | 0.52 (0.06) | 2.77 (0.41) | -13 | -13 | 0.49 | -20.7 (0.6) | 51.6 (4.2) |
| | 2017 | 39.4 (7.4) | 0.67 (0.08) | 2.60 (0.27) | -21 | 29 | 0 | -22.3 (0.6) | 62.0 (4.2) |
| 2-species | 2001 | 61.4 (12.7) | 0.60 (0.08) | 3.40 (0.75) | | | | -16.9 (1.1) | 25.9 (7.3) |
| (3) | 2011 | 53.3 (7.4) | 0.52 (0.07) | 2.72 (0.27) | -13 | -13 | 0.50 | -20.6 (5.7) | 50.6 (5.7) |
| | 2017 | 46.8 (4.9) | 0.46 (0.05) | 2.17 (0.30) | -12 | -12 | 0.51 | -22.1 (1.7) | 60.8 (11.5) |
| 3-species | 2001 | 61.4 (12.7) | 0.59 (0.08) | 3.60 (0.70) | | | | -16.8 (1.0) | 25.6 (6.7) |
| (3) | 2011 | 46.8 (12.7) | 0.53 (0.07) | 2.47 (0.59) | -24 | -10 | 0.70 | -22.0 (0.8) | 59.9 (5.1) |
| | 2017 | 46.9 (10.0) | 0.50 (0.09) | 2.38 (0.86) | 0 | -6 | 0 | -21.7 (1.2) | 58.3 (7.8) |
| 5-species | 2001 | 57.7 (10.3) | 0.59 (0.06) | 3.38 (0.63) | | | | -17.0 (0.9) | 26.8 (5.8) |
| (6) | 2011 | 52.4 (5.9) | 0.54 (0.04) | 2.81 (0.32) | -9 | -8 | 0.52 | -21.5 (0.9) | 56.3 (6.3) |
| | 2017 | 47.1 (8.5) | 0.45 (0.05) | 2.09 (0.36) | -10 | -17 | 0.38 | -22.3 (1.1) | 61.7 (7.5) |

### Supplementary Methods

*Aboveground tree carbon*

Aboveground tree biomass estimates were based on species- and diversity-specific allometric equations developed after harvesting and measuring 150 and 167 trees in the experiment in 2005 and 2017, respectively. A minimum of 9 trees per species and diversity treatment were harvested, encompassing the distribution of tree size. Diameter and height were recorded at the same time as tree harvesting. Biomass components were separated between trunk, branches, and leaves and weighed in the field. Samples of each component per individual tree were taken to a nearby laboratory where fresh weight was measured. Dry mass of all samples was then obtained after several days of drying at 105°C until constant mass. Average dry matter content (ratio of the sample dry mass to fresh mass) was calculated per species and tree size class and used to determine the dry biomass of branches and trunk (dry matter content × field fresh weight). Aboveground tree biomass was calculated as the sum of trunk and branch biomass (excluding leaves to focus on the more permanent C-components of the trees). The derived model was as follows (adjusted independently for the 2005 and 2017 datasets and for each species):

$$\ln(AGB_{t,d}) = a_d \times b_d \ln(X_{t,d}) + \varepsilon_d \,, \quad (1)$$

where AGB is the aboveground tree biomass, diversity level (d) is a three-level factor (1, 2/3, 5), and a and b are adjusted parameters. For each species and period, a set of different dependent variables (X) was tested, which included basal area at breast height (BA), diameter at 0.1m (basal diameter, Dbasal), and the products of BA and height (BaH). In case of multi-stemmed trees, BA is the sum of the basal area of all trunks. Model selection was based on AICc. Only one dependant variable was introduced at a time in the model, in order to keep meaningful degrees of freedom for model adjustment. Details and the best-fitting models are provided in Table S3. The best-fitting models were then combined with diameter and height inventories to estimate the ln(AGB) of all trees of the experiment in 2005 and 2016. We additionally estimated biomass in 2012, using the tree inventory conducted in 2012 and the best-fitting allometric equations calibrated in 2017.

The log-transformation of the data during the adjustment of allometric equations can lead to a bias in the final biomass estimation and uncorrected biomass estimates are theoretically expected to underestimate the real value. After the conversion of biomass from log to real unit, a correction was applied by multiplying the estimates by a correction factor CF, as suggested by Chave et al. (2005).

$$CF = \exp(\frac{RSE^2}{2}) \quad (2)$$

where RSE is the residual standard error. The uncertainty of biomass estimate due to the allometric models was provided; and calculated following Chave et al. (2004).

$$U = \sqrt{CF^2 - 1} \times < AGB > \quad (3)$$

where <AGB> is above-ground biomass estimate, CF is correction factor and U is the standard deviation measuring the error due to the allometric model.

AGB was converted to aboveground tree C (AGC) with species-specific trunk C concentrations obtained from coring tree trunks in the vicinity of the experiment (Hc: 45.07 % C; Ls: 45.76 % C; Ae: 45.82 % C; Tr: 47.01 % C; Co: 47.39 % C) (Elias and Potvin, 2003).

Table S3 Parameters of the allometric equations calibrated on observations from two field campaigns. Only the dependent variable and the inclusion of the interaction effect of tree diversity on parameters a and b vary among equations. Variables BA, Dbasal, BaH, Biomass (Y), are given in $cm^2$, cm, $cm^2$*m, kgDM, respectively.

| **Species** | **year** | **X** | **a main** | **a diversity=1** | **a diversity=3** | **a diversity=6** | **b main** | **b diversity=1** | **b diversity=3** | **b diversity=6** | **R²adj** |
|---|---|---|---|---|---|---|---|---|---|---|---|
| *Anacardium excelsum* | 2005 | Dbasal | | -5.49 | -5.51 | -5.48 | 3.31 | | | | 0.95 |
| | 2012 2016 | BA | | -1.60 | -1.43 | -1.39 | 1.09 | | | | 0.96 |
| *Cedrela odorata* | 2005 | Dbasal | | -5.36 | -0.09 | -5.20 | | 3.20 | 1.34 | 3.23 | 0.81 |
| | 2012 2016 | BA | | -2.62 | -2.623 | -2.622 | 1.25 | | | | 0.98 |
| *Hura crepitans* | 2005 | Dbasal | | -4.97 | -5.52 | -5.53 | 3.00 | | | | 0.87 |
| | 2012 2016 | BaH | | -2.89 | -2.96 | -3.06 | 0.89 | | | | 0.97 |
| *Luehea seemannii* | 2005 | Dbasal | | -4.99 | -5.27 | -5.00 | 3.06 | | | | 0.87 |
| | 2012 2016 | BA | | -0.65 | -0.99 | -2.34 | | 0.90 | 0.98 | 1.20 | 0.95 |
| *Tabebuia rosea* | 2005 | Dbasal | | -4.87 | -1.03 | -7.68 | | 2.98 | 1.45 | 4.19 | 0.77 |
| | 2012 2016 | BA | | -2.05 | -2.25 | -2.29 | 1.19 | | | | 0.97 |

#### *Coarse root carbon*

To estimate coarse root C (CRC) we relied on root:shoot ratios from the Sardinilla site based on two different root excavation campaigns. For CRC in 2005, we relied on root:shoot ratios developed in 2004 from the excavation of three-year-old trees, where ratios were obtained for Ls, Co, and Hc, and mean values were used for Ae and Tr (Coll et al., 2008). For CRC values in 2012 and 2016, we used species-specific root:shoot ratios developed in 2017 (16 year old trees), as described in Guillemot et al. (2020). The species-specific (but not diversity-specific) root:shoot ratios were then applied to each tree and multiplied with its AGC to obtain CRC estimates of all trees in the experiment.

#### *Coarse woody debris carbon*

All visible branches and stems fallen on the ground were collected annually in each plot and weighted to obtain a measure of coarse woody debris (CWD) biomass. We used samples collected in 2007, 2011 and 2016 which were collected at the end of the dry season (around March), except for 2007 where the collection lasted from July to August. CWD was put back after weighing. CWD biomass was converted to C (hereafter CWDC) using the species-specific trunk C concentration detailed above. For mixtures, we used the mean C concentration of the constituent species.

#### *Herbaceous carbon*

To estimate the biomass of the herbaceous vegetation, each plot was divided into four subplots of equal size and in each of these 88 subplots, herbaceous vegetation was cut to ground level in one randomly positioned, non-permanent, quadrat (0.5 $m^2$) twice a year, at the beginning and end of the wet season (May/June and November/December) as detailed in Potvin et al. (2011). Here, we use data for 2006, 2012, and 2016. The green and dry biomass collected in each quadrat was separated, dried, and weighed. The dry and green biomass within each of the four subplots of a given plot was summed and scaled to 1 $m^2$, considering individual plot size. To estimate the C concentration of herbaceous biomass, herbaceous vegetation samples were collected under 60 focal trees distributed according to species and diversity. The samples were separated into (1) legumes and (2) grasses / non-leguminous herbs and the C concentration was measured with an elemental analyser (Vario EL III, Elementar Analysensysteme, Hanau, Germany). There was no significant diversity effect on C concentration and the mean proportion of legumes versus grasses / non-leguminous herbs was ~50% across plots. We therefore used the mean

C concentration (42.72%) of legumes (44.78%) and grasses / non-leguminous herbs (40.66%) to convert herbaceous biomass to herbaceous C (hereafter herbaceous C).

*Leaf litter carbon*

Leaf litter was collected in 3–6 litter traps of 1 $m^2$ per plot every two weeks, with trap numbers varying with the number of tree species per plot. Litter collection lasted only 12 weeks in 2005 but 12 months in the other years. Here, we use data for 2005, 2012, and 2016. In 2005, we placed 6 litter traps in each of the 5-species plots and 3 litter traps in each of the 2-3 species plots and the monocultures. From 2012 onward, we placed 5 litter traps in each of the 5-species plots, 4 litter traps in each monoculture plot, 6 litter traps in each of the 3 species plots, and 4 litter traps in each of the 2-species plots. This later design was adopted to ensure an equal sample size when controlling for the identity of the tree species besides which a litter trap was positioned. As explained in Scherer-Lorenzen et al. (2005), each trap was positioned 1 m away from a tree of each species present in each plot. After litter bags were collected, the litter was dried at a constant temperature of 70ºC in the drying room of the Smithsonian Tropical Research Institute. Leaf litter production was calculated by dividing total dry biomass in grams from each trap by the number of days between two litter collection dates to determine the rate of litter fall per day per $m^2$. Litter biomass production was converted into litter C production (hereafter 'litter C') by using plot- and species-specific C concentrations measured from dry season litter (Scherer-Lorenzen et al., 2007). For mixtures, we used the plot-specific mean C concentration of the constituent species.

*Soil organic carbon*

In June 2001, June 2011, and April 2017, four soil cores were collected from each plot, one from each quadrat, measuring 5 cm in diameter, 10 cm in depth with the surface litter removed before sampling. The soil cores were dried and used to determine dry bulk density and analyzed for SOC concentration (%), and $\delta^{13}$C values at the Stable Isotope Laboratories at GEOTOP, UQAM, Montreal (2001) and UC Davis (2011 and 2017) (https://www.geotop.ca/fr/laboratoires/isotopes-stables-UQAM and https://stableisotopefacility.ucdavis.edu/, respectively). The bulk density and C concentration data were used to calculate the SOC stock (kg $m^{-2}$) and its $C_3$- and $C_4$-derived fractions. To understand the SOC stocks and fluxes we report it is important to take the land-use history of the examined site into account. As detailed briefly in the methods, the pasture that existed prior to the establishment of the Sardinilla planted forest was dominated by $C_4$ grasses which are thus the main contributor to the $SOC_4$ stocks we observe. Moreover, this pasture which existed prior to the planted forest, was established on a former rainforest site (Moore et al., 2018) and hence some of the rainforest-derived $SOC_3$ might still be left in the soil and contribute to the $SOC_3$ stocks we are reporting in Fig. 1. However, given that we calculate the changes in $C_3$-derived SOC, the former rainforest-derived $SOC_3$ does not play a role for the fluxes we examine (Table 2, Figures 3 and 5). More details on the sampling method and soil analysis can be found in Moore et al. (2018), which focused exclusively on SOC in the Sardinilla experiment.

*Leaf litter decomposition*

Leaf litter decomposition, hereafter litter decomposition, experiments were carried out in 2005, 2012, and 2017. In 2005, during a 116-day period during the wet season, 216 1-mm-mesh nylon bags of 25 x 25 cm in size were filled with 10 g of dry litter collected from litter traps and pinned down on the soil surface in the centre of the plots. The proportion of species within each bag represented the abundance of litter production species within each mixture. Litter bags were collected after 28, 56, 84, and 116 days, and subsequently dried and weighed (Scherer-Lorenzen et al., 2007). In 2012, a total of 12 g dry weight of air-dried (40°C) litter collected bi-weekly from traps located in monocultures and mixtures was placed carefully in identical decomposition bags as in 2005. Litter with signs of decomposition, herbivory or other damages was discarded. For species mixtures, equal proportions of litter from each species was used. Litter bags were installed with plot-specific litter. Monocultures and mixtures with three plant species mixtures, in which Ca was initially planted, were excluded. Litter bags were collected three times at 28, 69–70, and 100–103 days in the wet season. Mass loss at each collection was determined by cleaning the remaining litter carefully using a small paintbrush and distilled water and subsequently, oven-drying the samples at 65°C for four days. Initial dry weight values at 40°C were converted to dry mass at 65°C based on conversion factors calculated for each species (n = 5 per species). In 2017, litter bags were collected three times at 35–42, 68–75, and between 105–112 days in the wet season following the same procedure as in 2012. Litter bags from one plot were always collected on the same day. To ensure comparability of the decomposition rates between three examined periods, we only included data from plots which were measured in each period in our analysis: n = 5 monocultures (one plot for each species), n = 3 three-species mixtures, and n = 3 five-species mixtures.

*Root decomposition*

Root material from ten-year-old trees for the five tree species was manually excavated in non-experimental plots present in Sardinilla. Decomposition bags (10 cm × 20 cm) were made out of 2 mm nylon mesh and 5 g dry weight (air dried 40 °C for four days) of 4th and 5th root orders were placed into each bag (Guerrero-Ramírez et al., 2016). For species mixtures, equal proportions of root from each species were used. Four decomposition bags were inserted at a depth of 20 cm in a diagonal position and separated from the next bag by a distance of 40 cm in the centre of each experimental plot (52 decomposition bags). Root bags were installed in September 2011 in the ten monocultures and in three five-species mixture plots (n =13) and collected four times at 50, 195, 310, and 488 days. Mass loss at each collection was determined by washing roots and oven-drying samples at 65 °C for four days. Initial dry weight values at 40 °C were converted to dry mass at 65 °C based on conversion factors calculated for each species.

*Soil microbial biomass*

Soil microbial biomass ($C_{mic}$) was assessed in 2013 and 2017. $C_{mic}$ was measured as substrate-induced respiration, that is, the respiratory response of microorganisms to glucose addition (Anderson and Domsch, 1978) from ~5 g of sieved (2 mm), fresh soil. To saturate catabolic microbial enzymes, 8 µg glucose $g^{-1}$ soil dry weight was added as an aqueous solution to the soil samples. Microbial biomass (µg $C_{mic}$ $g^{-1}$ dry weight soil) was calculated as 38 × MIRR (µl $O_2$ $h^{-1}$ $g^{-1}$ dry soil) according to (Beck et al., 1997). MIRR, the maximum initial respiratory response, was assessed as the lowest respiration in the first ten hours – a period where microbial growth has not started. All measurements were conducted at 20°C in an air-conditioned laboratory using the same analytical devices (RMS Schuller, Darmstadt, Germany) (Cesarz et al., 2022).

*Soil respiration*

Soil respiration was measured with a portable infrared gas analyser (EGM-4, PP Systems, Amesbury, MA, USA) equipped with a soil respiration chamber (SRC-1, PP Systems, Amesbury, MA, USA). Attached to the soil respiration chamber was a PVC tube (10 cm in diameter, 10 cm long). Soil respiration was measured on the surface of the forest floor by gently pressing the chamber down on the forest floor during measurements to avoid cutting fine roots. The $CO_2$ concentration was measured every 5 sec over 90-120 sec between 09:00 and 15:00 LT (local time), and the change in $CO_2$ concentration over time was recorded. Soil respiration was calculated as follows:

$$\text{Soil respiration } (\mu\text{mol m}^{-2}\text{ s}^{-1}) = \left(\frac{\Delta \mathrm{CO_2}}{\Delta \mathrm{t}}\right) \times \frac{(P \times \mathrm{V})}{(R \times \mathrm{T} \times \mathrm{A})} \quad (4)$$

where $\Delta CO_2$ is the change in $CO_2$ concentration (µmol $mol^{-1}$) over time (t), calculated as the slope of the linear regression; P is the atmospheric pressure (Pa), V is the volume of the chamber including PVC tube ($m^3$), R is the universal gas constant (8.314 $m^3$ Pa $K^{-1}$ $mol^{-1}$), T is the temperature (K), and A is the surface area of ground covered by the chamber (0.007854 $m^2$). In each plot, soil respiration was measured at six to eight randomly chosen locations. Measurements were conducted in the wet season (late July and early August 2017).

*Canopy opening*

We measured canopy opening by taking four hemispheric photos per plot at 1 m above ground in October or November when trees were fully leaved-out following Sapijanskas et al. (2014) (Appendix b). In 2006 and 2012 we used a Nikon Coolpix 990 camera with the Nikkor E8 7.2 mm fish-eye lens. In 2016 the photos were taken by a Canon Eos Rebel XS camera and a Sigma Ex circular fisheye 4.5 mm lens. As detailed in Sapijanskas et al. (2014), the photos were analysed by the Gap Light Analyser (GLA) program (Frazer et al., 1999) to extract canopy structure from hemispheric photos. Canopy opening (a gap fraction in %) can be considered a measure of canopy space filling, which may mediate tree diversity effects on C stocks and fluxes (see Supplementary Table S1).

**Supplementary Analysis**

While we recognize that the small sample size limits the ability to test for normality, we nevertheless performed both Shapiro-Wilk normality test and examined the Q-Q plot to assert the extent to which data departed from normality. These tests were performed independently for each time period, showing no departure from normality for 45 of the 58 combinations of variables by time period. Departure from normality for the 13 combinations for which the Shapiro-Wilk test was significant was largely due to the presence of extreme values for some observations. Each one of these was verified and only in the case of CWDC did we decide to remove two points and performed the analyses with a reduced data set. In the other cases, the extreme values tended to be related to plots that either over- or under-performed because of either their species composition or the topography of the plots. When a plausible biological explanation was found, we decided to keep the data even if the normality assumption was not perfectly met.
